# Modeling neural coding in the auditory midbrain with high resolution and accuracy

**DOI:** 10.1101/2024.06.17.599294

**Authors:** Fotios Drakopoulos, Lloyd Pellatt, Shievanie Sabesan, Yiqing Xia, Andreas Fragner, Nicholas A Lesica

**Affiliations:** Ear Institute, University College London, United Kingdom; Perceptual Technologies, United Kingdom

## Abstract

Computational models of auditory processing can be valuable tools for research and technology development. Models of the cochlea are highly accurate and widely used, but models of the auditory brain lag far behind in both performance and penetration. Here, we present ICNet, a model that provides accurate simulation of neural dynamics in the inferior colliculus across a wide range of sounds, including near-perfect simulation of responses to speech. We developed ICNet using deep learning and large-scale intracranial recordings from gerbils, addressing three key modeling challenges that are common across all sensory systems: capturing the full statistical complexity of neuronal response patterns; accounting for physiological and experimental non-stationarity; and extracting features of sensory processing that are shared across different brains. ICNet can be used to simulate activity from thousands of neural units or to provide a compact representation of central auditory processing through its latent dynamics, facilitating a wide range of hearing and audio applications.

## Introduction

Computational models of sensory systems are widely used in fundamental research. They provide a platform for synthesizing existing knowledge and allow for generation and initial testing of hypotheses without the need for in vivo experiments. They can also facilitate the development of technologies that seek to approximate natural sensory function in artificial systems. But the utility of a model for a particular application depends on the degree to which it provides an accurate simulation of the true biological system at the relevant spatial and temporal scales. Numerous high-resolution models of the cochlea have been developed that faithfully capture its mechanical and electrical properties [1], with widespread uptake across academia and industry. Models of auditory processing in the brain, on the other hand, are much less accurate, with even the best missing out on a significant fraction of the explainable variance in sub-cortical and cortical neural activity [2, 3].

Models of the cochlea have typically been designed by hand to mimic the transformations performed by the biological system. This approach works well for the cochlea because it contains only a few stages with relatively homogeneous elements. The brain, however, with its many interconnected and diverse neural networks, is too complex for such an approach. Hand-designed models of the auditory brain can be effective in describing specific features of responses to a limited class of sounds [4–7], but more general models of the brain require the use of “black box” frameworks with parameters estimated from neural recordings. To facilitate model fitting from limited data, the allowable transformations are typically constrained to be either linear or low-order non-linear [2]. These constraints aid interpretability, but such models lack the capacity to capture the full complexity of auditory processing and, thus, are not accurate enough for many applications. Advances in deep learning have created a new opportunity to develop accurate models of sensory processing in the brain, making it feasible to fit models that capture arbitrary non-linear transformations directly from data. Recent studies have shown that deep neural network (DNN) models of the sensory periphery can be highly accurate. DNNs perform similarly to biophysical models of the cochlea in predicting a wide range of auditory phenomena [8–11] and, beyond the auditory system, models of the retina provide accurate predictions of ganglion cell responses to natural scenes [12]. Initial attempts to build DNN models of sensory processing in the brain have also produced impressive results. Models of primary visual cortex (V1) responses to natural images explained 50-90% of the explainable variance in recorded activity [13–15], while models of V4 explained 90% [16]. DNN models of primary auditory cortex (A1) responses to natural sounds perform similarly well, explaining 60-70% of the explainable variance in recorded activity [17, 18]. But these models reproduce only low-resolution measurements of activity (calcium transients [13, 15], spike counts over large time windows [14, 16], field potentials [18] or fMRI voxels [17]), ignoring potentially important information in the neural code that is present at fine spatial and temporal scales. One recent attempt to use DNNs to model single- and multi-unit spiking in auditory cortex with a temporal resolution of 10 ms succeeded in capturing approximately half of the explainable variance [19].

For a model of neural coding in the early auditory pathway, both high spatial and temporal resolution are essential [20]. We recently developed a DNN model of the inferior colliculus (IC), the midbrain hub of the central auditory pathway, to simulate neural response patterns with millisecond precision [21]. We trained the model on multi-unit activity (MUA) recorded across the extent of the IC, with the goal of identifying and capturing latent neural dynamics that reflect the fundamental computations that are common to all brains, rather than the idiosyncrasies of particular neurons. This model was effective in predicting mean responses to a limited set of speech sounds. However, to bring its performance up to the level required for it to serve as a useful general model of central auditory processing or as a valid in silico surrogate for the IC in auditory neuroscience, there are three key challenges that must be addressed. The first is to capture the full statistical complexity of neural activity. Models of neural coding typically rely on regression frameworks with assumed distributions (e.g., Poisson) that constrain the relationship between response strength and reliability [2]. While these assumed distributions might be valid for some higher-level brain areas, they are a poor match for early sensory pathways where strength and reliability exhibit complex interactions.

The second challenge is to address non-stationarity. Even in the absence of behavioral modulation, the mapping between sensory input and recorded neural activity can change over time for both physiological reasons (e.g., changes in brain state) and experimental reasons (e.g., movement of the recording electrode within the brain). Because training high-capacity models requires the use of large datasets recorded over long periods of time, non-stationarity is inevitable and must be explicitly accounted for. The final challenge is to overcome the idiosyncrasy of individual brains and the recordings made from them. While models must be trained on individual recordings, the final model of a sensory brain area should reflect only the processing that is common across all healthy individuals.

To address these challenges and derive an accurate and general model of early central auditory processing, we developed a novel encoder-decoder framework for modeling neural coding which we trained on large-scale recordings from the IC of anesthetized gerbils. The framework comprises a shared encoder that maps a sound waveform to a generic latent representation, followed by separate decoders that map the generic representation to the neural activity of individual brains. Each decoder also receives a second input that serves as a time stamp, which it uses to modulate the generic representation to account for non-stationarity specific to each brain. This allows the model to decouple the fundamental aspects of auditory processing from the effects of physiological degradation or electrode drift so that, once fully trained, it can generate outputs that reflect optimal recording conditions. The final model, which we call ICNet, can simulate neural activity across many brains in response to a wide range of sounds with high accuracy and fast execution, and its latent dynamics provide a compact description of auditory processing that can serve a variety of applications. ICNet can be used as a standalone model with outputs that reflect the baseline neural representation at the level of the IC, or as a foundation core for models that include additional aspects of auditory processing such as behavioral modulation or higher-level computations.

In the following sections, we describe the DNN-based framework that we used to develop ICNet and then evaluate its performance. We first describe the DNN components that we designed to address each of the three modeling challenges. We then present the final model and demonstrate how its encoder can be used as a front-end for automatic speech recognition. We establish the remarkable accuracy of the model by comparing neural responses simulated by ICNet with those recorded from the IC for a variety of sounds that were not part of training (including speech in quiet and in noise, moving ripples, and music at different intensities). Finally, we demonstrate that ICNet also captures fundamental neurophysiological phenomena and further establish its generality across new animals and sounds.

## Results

To collect the dataset required for model training, we recorded neural activity from the IC of nine normal-hearing gerbils, a common animal model for human hearing, using custom-designed electrode arrays with 512 channels (Fig. 1a) [21, 22]. We made the recordings under anesthesia, allowing us to present a wide variety of sounds to each animal across a range of intensities, including speech in quiet and in noise, processed speech, music, environmental sounds, moving ripples and tones that together totaled more than 10 hr in duration. We processed the recordings to extract MUA (threshold crossing counts on each recording channel in 1.3 ms time bins) [20, 21]. We included all recorded units in model training, but only those units with a signal correlation (correlation of responses across repeated trials of broadband noise) of 0.2 or higher are included in our model evaluation (494 ± 16.7 units from each animal, resulting in a total of 4446 units across all animals; see Supplementary Fig. 1).

**Fig. 1.**
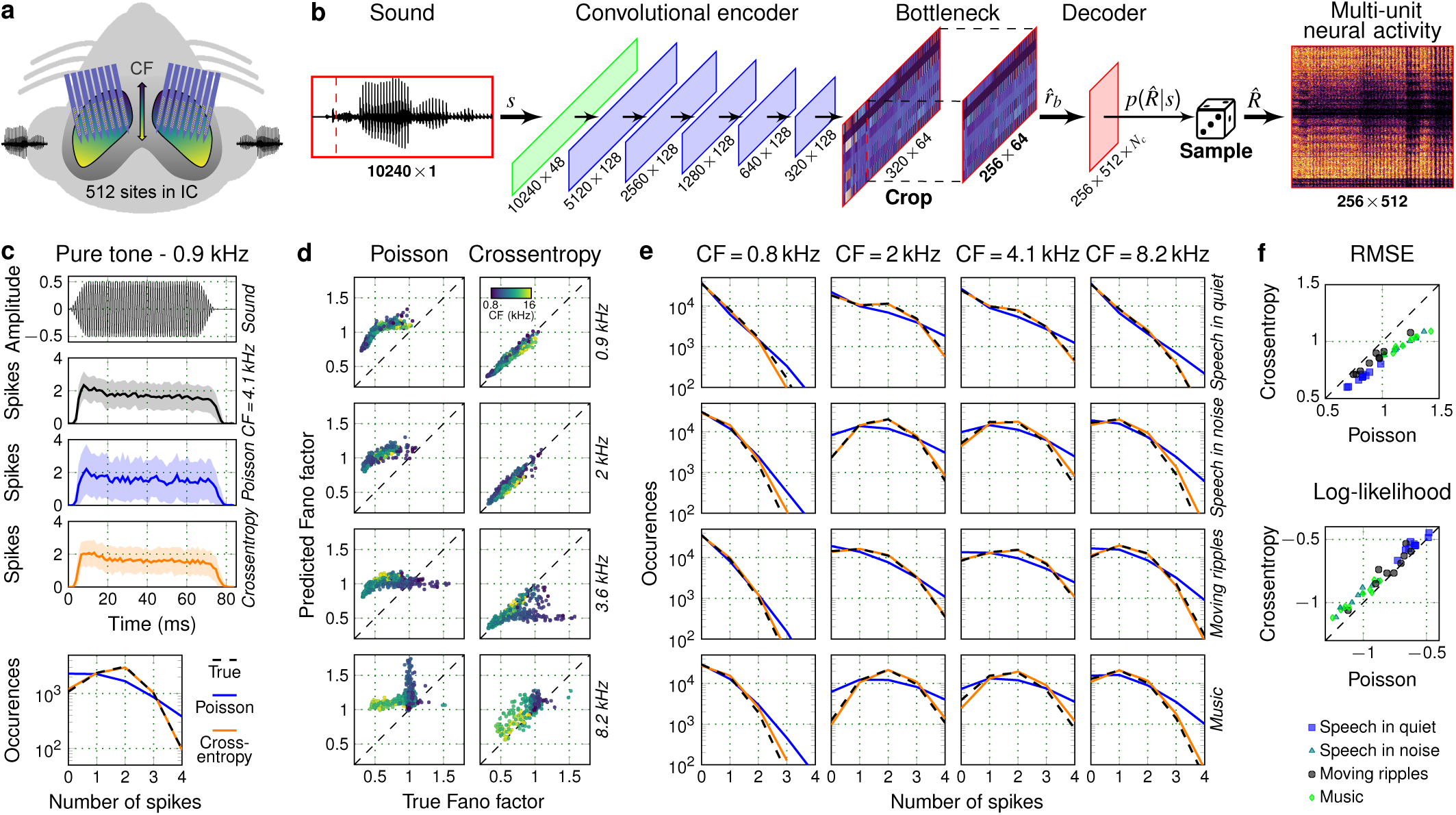
Modeling the statistical properties of neural activity. **a.** Schematic of the geometry of custom-designed electrode arrays for large-scale recordings in the gerbil inferior colliculus (IC). The recording sites were distributed across 8 electrode shanks and spanned a plane measuring 1.4 × 0.45 mm within each hemisphere. **b.** Schematic diagram of the deep neural network (DNN) architecture trained to map a sound waveform to neural activity for 512 units from the IC of a single animal. *N_c_* is the number of parameters in the probability distribution *p*(*R*^^^|*s*), which is 1 for Poisson models and 5 for crossentropy models. **c.** Peri-stimulus time histograms (PSTHs) of recorded and predicted neural activity for one unit (characteristic frequency (CF) of 4.1 kHz) in response to 128 presentations of a 0.9 kHz pure tone at 85 dB sound pressure level (SPL). Neural activity was simulated using either a Poisson or crossentropy model. The shaded regions indicate the standard deviation across the 128 trials. The bottom panel shows the distribution of MUA counts across trials for the responses in the PSTH plots, averaged across time bins during the presentation of the tone. **d.** Comparison of the Fano factor (variance-to-mean ratio) between recorded and predicted neural activity (each point corresponds to one of 483 units from one animal) for the Poisson and crossentropy models (columns) in response to 4 tones (rows). Points are colored based on the CF of the corresponding unit, with brighter colors indicating higher CFs. For further discussion of model accuracy for narrowband sounds, see Supplementary Fig. 2. **e.** Comparison of MUA count distributions for recorded and predicted neural activity for 4 units (columns) in response to 4 sounds (rows). **f.** Performance comparison across 9 animals and 4 sounds between Poisson and crossentropy models. Each symbol represents the average across all time bins and units for one animal in response to one sound.

We trained DNN models to simulate neural coding, i.e., to take as input each presented sound waveform and produce as output the corresponding MUA spiking patterns recorded from all units in each individual animal. These models use a cascade of convolutional layers to encode the sound waveform *s* to a latent representation *r*^*_b_* (bottleneck), which is then passed to a simple linear decoder to predict the response *R*^^^ for the recorded units (Fig. 1b). The output of the decoder for each unit in each time bin defines a probability distribution *p*(*R*^^^|*s*), from which we can then sample to simulate the activity *R*^^^ of each unit across time with millisecond precision. Once trained, the DNN models can be used to simulate the neural activity of all recorded units for any sound, whether it was presented during neural recordings or not.

### DNNs can capture the statistics of neural response patterns

We first trained a set of baseline models to simulate activity across all units for each animal through Poisson regression [21] (*N_c_* = 1 in the decoder layer of Fig. 1b to produce the Poisson rate parameter *λ*). The Poisson assumption is common in models of neural coding. It provides a theoretical basis for the optimal estimation of model parameters via classical statistical methods, and it is a reasonable approximation of the true distribution of activity in some brain areas. But activity in sub-cortical areas like the IC is typically underdispersed, i.e., it is much more reliable across trials than is expected for a Poisson process [23, 24]. This phenomenon is illustrated in Fig. 1c, which compares the peri-stimulus time histograms (PSTHs) of recorded and simulated neural responses for an example IC unit to 128 trials of a pure tone. The Poisson model captured the mean activity across trials well (indicated by the solid lines), but overestimated the variability across trials (indicated by the shaded areas).

Fortunately, DNNs can be trained to predict full activity distributions without any assumptions by framing the problem as classification rather than regression [25]. We trained the same DNN architecture for each animal with a crossentropy loss and categorical distributions with *N_c_* = 5 discrete classes (corresponding to the probability of each unit having a count of 0 to 4 within a given 1.3 ms time bin). The crossentropy model accurately captured both the mean activity and the variability across trials for the example unit (Fig. 1c). For all time bins within the duration of the tone, the Fano factor (ratio of variance to mean) was 0.55 for the true response, 0.54 for crossentropy model, and 1.16 for the Poisson model. (Note that because the rate of the Poisson model varies across time bins, the overall Fano factor of its activity does not need to be exactly 1.) For all units from one example animal in response to four different tone frequencies, the Fano factor for the crossentropy model was much closer to the true value than the Fano factor for the Poisson model (Fig. 1d; the average absolute error was 0.38 for the Poisson model and 0.07 for the crossentropy model).

To compare the overall performance of the Poisson and crossentropy models, we simulated responses for all animals to four different sounds: speech in quiet, speech in noise, moving ripples, and music. As shown in Fig. 1e for four example units, the crossentropy model accurately predicted the full MUA count distributions for each sound. We compared the overall performance of the models for each animal and sound using two metrics (averaged across all time bins and units): the root-mean-square error (RMSE) between the recorded and predicted activity, and the logarithm of the probability (log-likelihood) of observing the recorded activity from the activity distribution inferred by the model. We found consistent improvements for the crossentropy models across all animals and sounds (Fig. 1f; improvement of 15.98% [15.88, 16.02] for RMSE (median and 95% CIs over bootstrap samples) and 8.13% [8.04, 8.16] for log-likelihood). These results demonstrate that characterizing the full distributions of spiking activity for each neural unit is feasible and improves the accuracy of simulated neural coding in the IC.

### DNNs can account for non-stationarity in neural recordings

Capturing the full non-linearity of auditory processing requires the use of high-capacity models, which in turn requires large datasets recorded over long periods of time. Non-stationarity in such recordings is inevitable, as the physiological state of the animal or the electrode position can vary over the course of the recording, altering the mapping from sound to neural activity expressed by individual units. Fig. 2a illustrates this for two of our recordings, where neural activity in response to the same speech sound was either stable (stationary) or variable (non-stationary) over the 10 hours of recording. While there is an overall trend of decreased activity in the second example, the non-stationarity can vary widely across individual units (Fig. 2b), creating a challenging modeling problem.

**Fig. 2.**
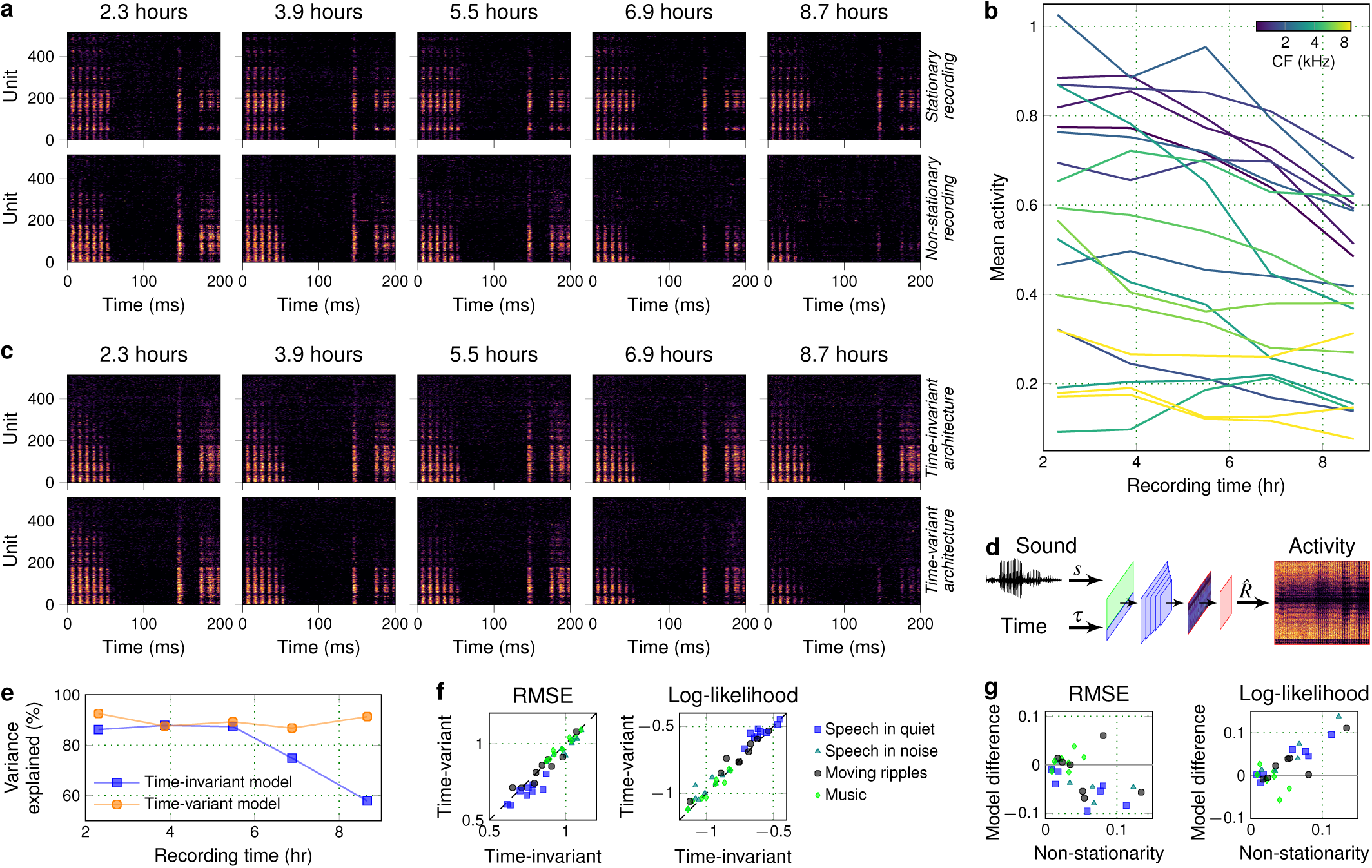
Accounting for non-stationarity in neural recordings. **a.** Example neural activity in response to the same sound (speech at 60 dB SPL) presented at five different times within two individual recordings. Units were ordered based on their CFs. Brighter colors indicate higher activity. **b.** The mean activity (per bin) of 20 randomly selected units from the non-stationary recording shown in **a**. **c.** Predicted neural activity for the non-stationary recording in **a** using the time-invariant and the time-variant models. **d.** Schematic diagram of the time-variant DNN architecture. **e.** Median predictive power of the time-invariant and time-variant models across all units at five different times during the non-stationary recording shown in **a**. Predictive power was computed from the simulated and recorded neural responses as the fraction of explainable variance explained for a 30-s speech segment presented at 60 dB SPL. **f.** Performance comparison across 9 animals and 4 sounds between time-variant and time-invariant DNN models. Each symbol represents the average across all time bins and units for one animal in response to one sound. **g.** Performance difference between models (time-variant − time-invariant) as a function of recording non-stationarity across 7 animals and 4 sounds.

A standard encoder-decoder model can only capture a time-invariant encoding that will reflect the average mapping from sound to neural activity over a large dataset. To overcome this limitation, we designed a time-variant version of the crossentropy model that takes an additional input which acts as a time stamp, reflecting the time during recording at which the sound was presented (Fig. 2d). The interactions between the two inputs within the encoder enable the model to alter its sound encoding as needed to account for non-stationarity. After the model is trained, neural activity can be simulated to reflect the sound encoding at specific time points during the recording (by providing the appropriate additional inputs), or the simulation can made stationary by keeping the additional input constant across all sounds.

As a first test, we trained time-invariant and time-variant models to predict the neural activity of the example non-stationary recording and evaluated their performance in predicting the responses shown in Fig. 2a. The time-invariant model, by construction, predicts the same responses each time a sound is presented. But the time-variant model learned to utilize the additional input to capture the changes over the course of the recording, predicting strong responses at early recording times and weak responses at late recording times (Fig. 2c) while capturing non-uniform and non-monotonic changes in activity across units (Supplementary Fig. 3). We quantified the differences in model performance for this example by computing the fraction of explainable variance in the recorded activity (i.e., the covariance across repeated trials) that each model explained across all units for each of the five speech presentations. The results demonstrate that the time-variant model can capture the changes in the mapping from sound to neural activity over time, consistently explaining close to 90% of the explainable variance across the entire recording (Fig. 2e), while the performance of the time-invariant model was much more variable and worse overall.

Across all animals and sounds, we found that the time-variant models improved performance by an average of 2.67% [2.60, 2.69] for RMSE and by 3.22% [3.11, 3.26] for log-likelihood (Fig. 2f). But we did not expect performance improvements across all animals, as the time-variant model should only achieve better predictions for recordings that are non-stationary. We categorized the recordings for which we had repeated trials of the same sounds presented at different times in the recording (seven out of nine animals) based on their non-stationarity. We measured non-stationarity as the absolute difference of the covariance of neural activity between two successive trials and two trials recorded several hours apart. As expected, we found that the benefit of the time-variant model increased with non-stationarity for both the RMSE and the log-likelihood measures (Fig. 2f), approaching 10% improvement on RMSE and 15% improvement on log-likelihood for the most non-stationary recordings. These results demonstrate that time-variant DNNs can flexibly model non-stationarity, improving performance for recordings that are unstable without compromising performance for stable recordings.

### Multi-branch DNNs can capture shared latent dynamics across animals

Our goal was not to model any individual animal, but rather to develop a generic model of the IC that describes the features of sound encoding that are common across all animals. Recordings from any individual animal will inevitably contain idiosyncrasies that reflect variations in anatomy or physiology, or in the position of the recording electrode within the IC. These idiosyncrasies may be critical to preserve when developing models for applications such as hearing aid development, which requires sound processing to be unique to each individual hearing loss profile. But for applications that require a generic model of normal auditory processing, architectures that can extract what is common across recordings from different individuals are more suitable.

We have previously demonstrated that the acoustic information in IC activity is restricted to a low-dimensional subspace which is shared across animals with the same hearing status [21]. This result suggests that it should be possible for a DNN with shared latent dynamics to predict neural activity for any normal-hearing animal. We thus designed a multi-branch DNN architecture comprising a shared encoder and separate decoder branches that are used to simulate neural activity for each of nine individual animals (Fig. 3a). To simulate activity for all animals with a shared encoder, we also had to modify the way in which the additional time input was used so that the non-stationarity of each individual recording could be accounted for and could be decoupled from the shared sound encoding. We introduced animal-specific time inputs at the level of the decoder and combined each time input with the bottleneck output, allowing for a unique time-variant mapping from the generic latent representation to the neural activity for each animal. We considered mappings of varying complexity for the time inputs, but found that a simple non-linear projection was sufficient for our purposes (Supplementary Fig. 3). After training, the multi-branch model, which we call ICNet, can simulate neural activity for nine individual animals in response to any sound in a manner that reflects the state of each animal at any given time.

**Fig. 3.**
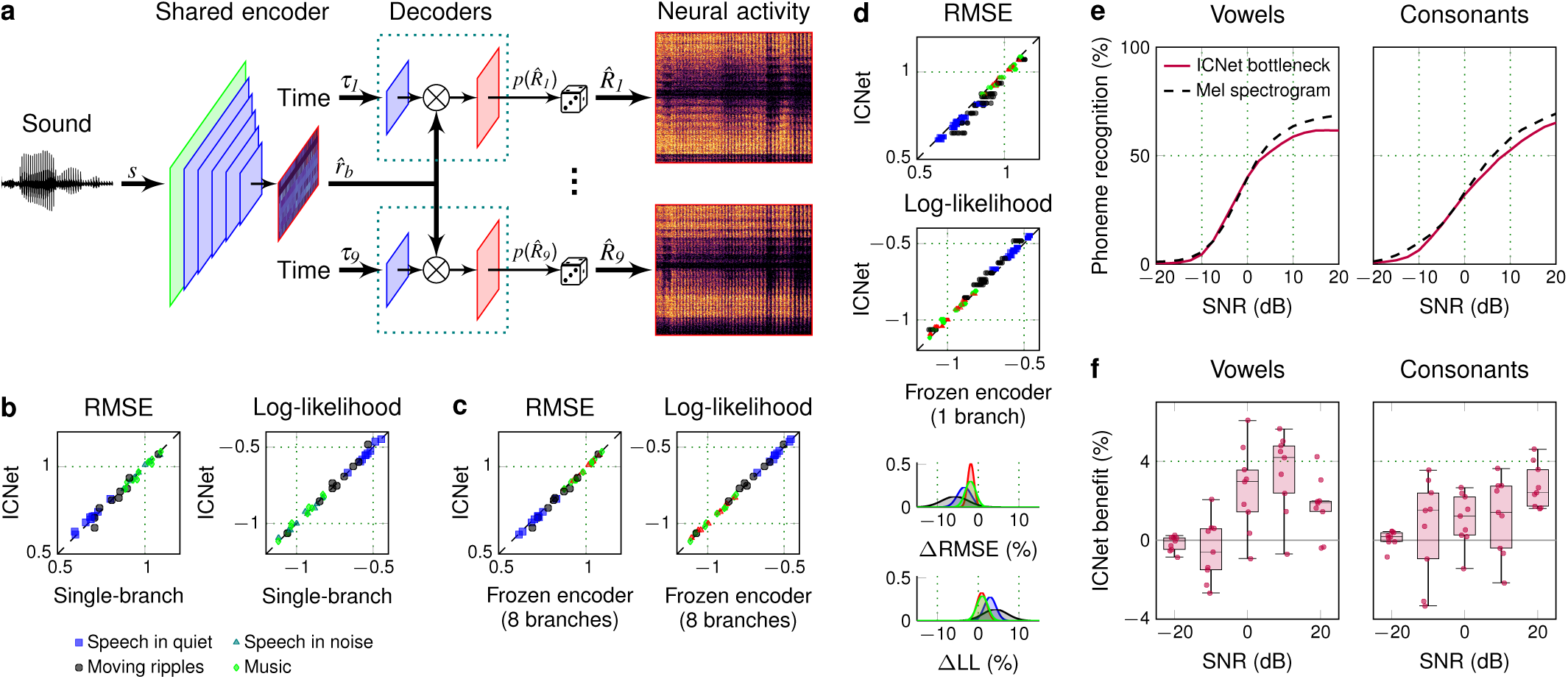
Capturing shared sound encoding across animals. **a.** Schematic diagram of the ICNet model trained to map a sound waveform to the neural activity of 9 animals. **b.** Performance comparison for 9 animals between ICNet and the corresponding single-branch models. Each symbol represents the average across all time bins and units for one animal in response to one sound. **c.** Cross-validation comparison between ICNet and individual eight-branch models with frozen encoders and decoders trained to predict the ninth animal. The similar performance across animals and sounds demonstrates that multi-branch models learn generic latent dynamics that can be transferred to other animals. **d.** Cross-validation comparison between ICNet and individual single-branch models with frozen encoders and decoders trained to predict the remaining eight animals. Distributions in the bottom show the spread of ICNet benefit for each sound compared to all 72 frozen-encoder models. The decrease in performance across animals and sounds demonstrates that single-branch models cannot fully describe the sound encoding of other animals. **e.** Vowel and consonant recognition performance of an automatic-speech-recognition (ASR) model trained to predict phonemes from either the ICNet bottleneck response *r*^*_b_* or a Mel spectrogram. Phoneme recognition was computed for sentences presented at 65 dB SPL and mixed with speech-shaped noise at SNRs between -20 and 20 dB. **f.** Benefit in recognition performance for the ASR model trained with the ICNet bottleneck response as input, compared to ASR models trained with the bottleneck responses of the 9 single-branch models.

We first evaluated ICNet by simulating neural activity in response to four sounds (with the time input matched to the time at which each sound was presented to each animal). ICNet matched the performance of the corresponding single-branch models for all nine animals (Fig. 3b; average RMSE increase of 0.04% [0.01, 0.06] and log-likelihood decrease of 0.39% [0.34, 0.41] compared to time-variant models trained to predict the activity of a single animal). To establish that ICNet encoded neural dynamics that were shared across all animals (rather than the idiosyncrasies of each), we performed a cross-validation experiment. We held out each of the nine individual animals in turn while training an eight-branch model to predict the neural activity of the remaining eight animals. We then froze the encoder of each eight-branch model and trained a new decoder that used the output of the frozen encoder to predict the neural activity of the held-out animal.

As shown in Fig. 3c, the models that used a frozen encoder and an optimized decoder performed similarly to the original ICNet (1.19% [1.16, 1.20] decrease of RMSE and 0.41% [0.38, 0.42] increase of log-likelihood for ICNet compared to the frozen-encoder models), i.e., the performance for an individual animal was similar whether or not that animal was included in the dataset used to train the encoder. This suggests that ICNet performs a generic encoding that can predict the neural activity of any normal-hearing animal. To further validate this, we made recordings from an additional three animals that were not included in the training of ICNet. We used these recordings to train either full single-branch models or new decoders to be used with the frozen ICNet encoder. We found that with only a few minutes of data, the performance of ICNet with the new decoders matched that of the single-branch models trained on the full dataset (Supplementary Fig. 4). We next reversed the cross-validation experiment to establish the benefit of our multi-branch DNN architecture, i.e., the degree to which the single-branch models captured the idiosyncrasies of each animal without fully capturing the neural dynamics that are shared across animals. We froze the encoder of each of the nine single-branch models and trained eight new decoders to predict the neural activity of the remaining animals. As shown in Fig. 3d, the difference in the performance of ICNet and these frozen-encoder single-branch models was small, but the frozen-encoder models were almost always worse, especially for speech in quiet and moving ripples (average RMSE decrease of 3.07% [3.05, 3.07] and log-likelihood increase of 2.01% [2.00, 2.02] for ICNet compared to the frozen-encoder models). Taken together, the results of these experiments establish that ICNet is a generic yet comprehensive model of sound encoding.

The design of ICNet with its complex encoder and simple decoders differs from that of standard encoder-decoder DNN models. Typically, a complex encoder is used to map the input to an abstract latent space, and a decoder of similar complexity is used to map the latent representation to the final output, allowing for a complex relationship between the low-dimensional latent representation and the high-dimensional output. The simple linear decoders in ICNet, on the other hand, ensure that the latent representation in the bottleneck is constrained to directly reflect the dynamics that underlie neural activity. And because the bottleneck is shared across the individual decoder branches (Fig. 3a), the bottleneck response *r*^*_b_*in ICNet reflects a general set of dynamics that is common to all animals and can be used on its own as a compact representation of central auditory processing.

As a demonstration, we used the ICNet sound encoder as an acoustic front-end and trained a simple automatic-speech-recognition (ASR) DNN to classify phonemes from the bottleneck response *r*^*_b_*. We found that the ASR system trained on the ICNet bottleneck response achieved similar performance to that of a baseline system trained with a Mel spectrogram input (Fig. 3e). We also found that training on the ICNet bottleneck response resulted in better performance than training the same ASR system with the bottleneck responses from the nine single-branch models. This difference was small overall, but increased at relatively high signal-to-noise ratios (SNRs) (Fig. 3f; average improvement of 1.24% [1.02, 1.76] for consonant recognition and 1.60% [1.25, 2.22] for vowel recognition across all SNRs and animals). These results demonstrate that the encoder of ICNet can serve in the same manner as widely used acoustic front-ends such as auditory filterbanks or cochlear models for applications where simulation of central (rather than peripheral) auditory processing is desired.

### ICNet is a highly accurate model of neural coding

Fig. 4a shows example segments of recorded activity and predictions from ICNet for four sounds, with each image showing the response of 1000 units (the units were selected to achieve a uniform distribution of characteristic frequencies (CFs) on a logarithmic scale, reflecting the tonotopic organization of the IC). ICNet’s simulations were nearly indistinguishable from the recordings across all units and all sounds. The full non-linearity of the mapping from sound to neural activity that is captured by ICNet is difficult to appreciate from visual inspection, but Fig. 4a illustrates some of its remarkable features. For example, the speech in quiet elicited the strongest activity even though it was presented at the lowest intensity of any of the sounds (60 dB SPL with energy concentrated at low frequencies), while the speech in noise, which was presented at a much higher intensity (85 dB SPL with a much broader range of frequencies), elicited significantly less activity. The response to the moving ripples was particularly complex: the sound was presented at high intensity (85 dB SPL) with all energy concentrated at high frequencies, yet it elicited relatively weak activity in units with high CFs and relatively strong activity in units with low CFs. This frequency transposition indicates a high degree of non-linearity that was captured well by ICNet.

**Fig. 4.**
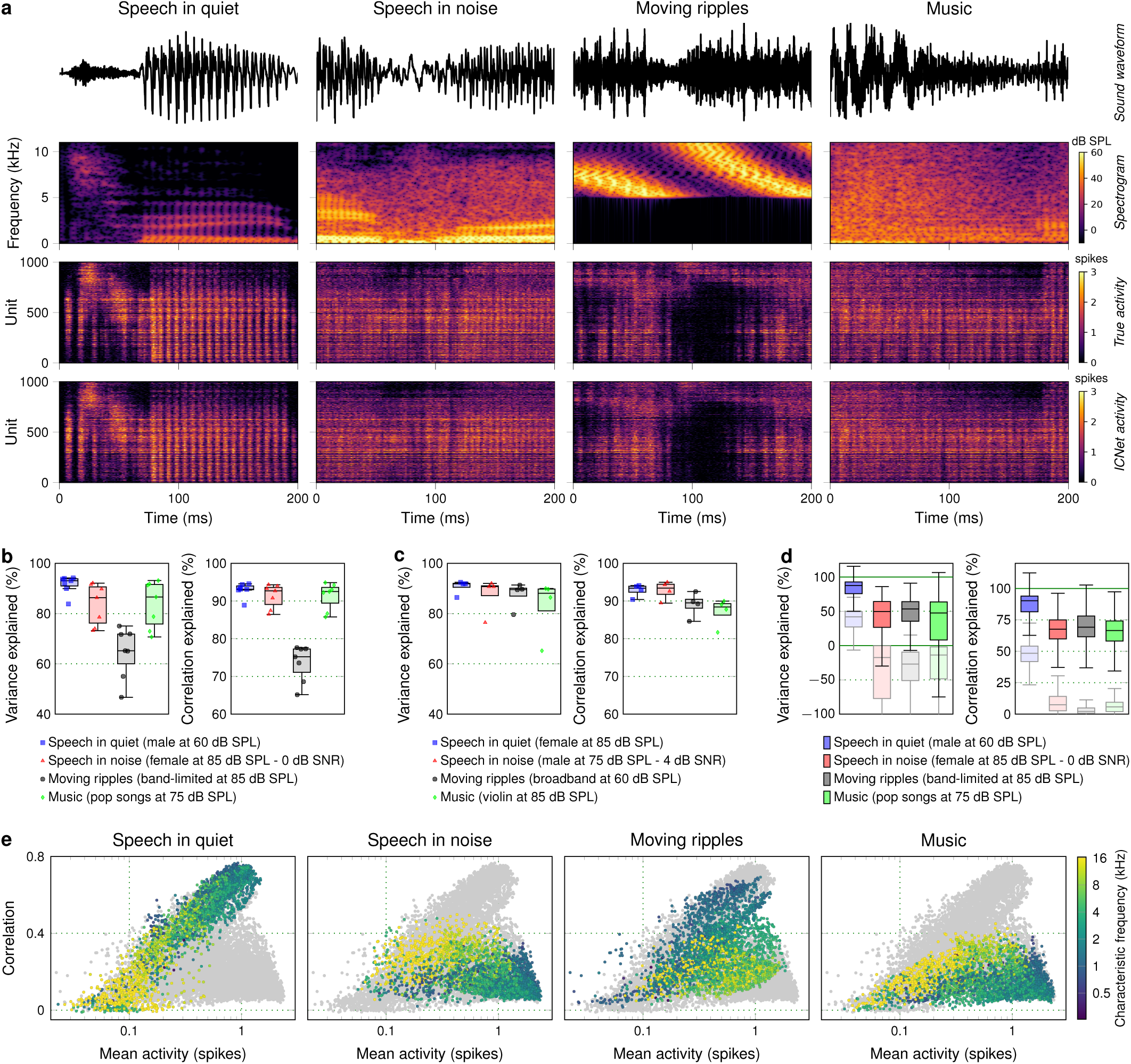
ICNet performance across sounds. **a.** Segments of recorded and ICNet predicted neural activity in response to 4 sounds. Each column shows the sound waveform and corresponding spectrogram, as well as the neural responses of 1000 units. Units were selected from all animals to achieve a uniform logarithmic spacing of CFs between 300 and 12000 Hz. Brighter colors indicate higher stimulus intensity for the spectrograms and higher activity for the neural responses. **b.** Predictive power of ICNet across 7 animals and 4 sounds, computed as the fraction of explainable variance and correlation explained by the model across all time bins and units of each animal. **c.** Predictive power of ICNet across 4 animals and 4 sounds, computed as the fraction of explainable variance and correlation explained by the model across all time bins and units of each animal. **d.** Predictive power of ICNet across 4 sounds and 3476 recorded units (from 7 animals), computed as the fraction of explainable variance and correlation explained by the model across all time bins of each unit. The shaded box plots show the predictive power of linear-nonlinear Poisson (LNP) models for the same units. **e.** Overall activity (mean spiking activity) versus reliability (correlation between two successive trials) for 3476 units from 7 animals. Each panel shows the results for all units in response to one sound, with each point colored based on the CF of the unit. The gray points in the background show the results for all four sounds and are the same in each subplot.

To provide an overall measure of the predictive power of ICNet, we used seven of the nine animals for which we had recordings of responses to the same sounds on successive trials. We computed the fractions of the explainable variance and correlation (the covariance and correlation of recorded responses on successive trials) that ICNet explained for sounds that were not included during training (measured after flattening the responses by collapsing across time and units for each animal; see Methods). We found that ICNet performed best for speech in quiet, explaining more than 90% of the explainable variance and correlation across all animals (Fig. 4b; 91.5% [91.4, 91.8] for variance and 92.9% [92.9, 93.1] for correlation).

Performance was also excellent for speech in noise (83.6% [83.3 84.3] for variance, 91.2% [91.1 91.6] for correlation) and for music (83.6% [83.3 84.3] for variance, 91.3% [91.2 91.7] for correlation), and decreased only for the band-limited moving ripples (64.4% [64.0, 65.2] for variance, 73.5% [73.3, 74.0] for correlation). To further test the generality of ICNet, we also measured performance for four additional sounds that were presented on successive trials for four of the nine animals: speech in quiet at a higher intensity, speech in noise at a higher signal-to-noise ratio (SNR), broadband moving ripples, and a solo violin recording. We found that ICNet again explained most of the explainable variance and correlation for all sounds (Fig. 4c; median values for variance and correlation between 88% and 93%).

The flattened metrics computed over all units are appropriate for assessing the overall predictive power of the model, but are dominated by the units with the most explainable variance or correlation (which are, presumably, the most important units to model accurately since their activity carries the most information about sound). When we instead examined ICNet’s performance across individual units, we found lower values overall and substantial variability across units and sounds (Fig. 4d). The lower values are a result of the difference in the way the flattened and unit-by-unit metrics account for the variation in mean activity across units for a given sound. This variation is fully accounted for by the flattened metrics but is either partly (for variance) or fully (for correlation) ignored when the metrics are computed unit by unit. This variation makes up a substantial portion of the explainable variance and correlation in the full neural activity and is well captured by the model, so the resulting performance values for the flattened metrics are higher than those for the individual units. ICNet outperformed standard single-layer linear-nonlinear Poisson (LNP) models of the same units by a wide margin (shaded box plots in Fig. 4d), especially for non-speech sounds. (Note that negative values for the variance explained by LNP models indicate that the variance of the error in the model prediction was larger than the explainable variance in the response (Eq *4*).)

The variability in performance across units could have many sources. From the responses in Fig. 4a, it is clear that units can vary in their activity depending on the match between the acoustic properties of a sound and the features to which a unit is responsive. Both overall activity (measured as mean count) and reliability (measured as the correlation of responses on successive trials) of individual units spanned a wide range that varied across sounds (Fig. 4e), with a clear dependence on CF. To determine whether any of these properties (overall activity, reliability, preferred frequency) could explain the variation in model performance across units, we measured their correlation with the fraction of explainable correlation explained across all sounds. We found only a weak correlation between overall activity and model performance (−0.12 [-0.14, -0.10] for 3406 units from seven animals, after excluding units with the top and bottom 1% of explainable correlation explained) and no correlation between CF and model performance (−0.02 [-0.03, 0]). But the correlation between reliability and model performance was much stronger (0.54 [0.53, 0.55]). This suggests that the reliability of neural responses to a given sound is a strong determinant of ICNet performance.

We also examined whether any of the errors in ICNet predictions were systematic, i.e, consistent for a particular sound class or response timescale (Supplementary Fig. 2). For our four primary evaluation sounds (speech in quiet, speech in noise, music and moving ripples), we found no evidence of systematic errors. The overall magnitude of predicted and recorded activity was well matched across all units and sounds, and the coherence spectrum between predicted and recorded activity mirrored that of the recorded activity across successive trials. ICNet predictions for narrowband sounds were also generally well matched to recorded responses, but there was a consistent overprediction of the responses of low-CF units to above-CF sounds at high intensities.

### ICNet captures fundamental neurophysiological phenomena

Having established that ICNet can provide accurate simulation of responses to complex sounds, we investigated whether it also captures more fundamental neurophysiological phenomena. We designed a new set of sounds based on those that have traditionally been used to characterize response properties of IC neurons, and presented the new sounds to two new animals along with an abridged set of our original training sounds (see Methods). We used the recorded training dataset to train new decoders for the two animals (keeping the ICNet encoder frozen) and then used the updated models to predict the responses to the new evaluation sounds. ICNet performance for the new sounds was excellent overall, with more than 80% of the explainable variance and correlation captured for all sounds, and again matched the performance of fully trained single-branch models with just a few minutes of training data (Supplementary Fig. 4).

We examined key response properties at the level of output units as well as in the bottleneck. Because the ICNet decoder is so simple, we expected to see direct manifestations of the phenomena underlying the response properties in the latent dynamics. We first examined simple properties of responses to tones. The frequency response areas (FRAs; heatmaps showing overall response magnitude as function of tone frequency and intensity) of ICNet output units were well matched to those derived from the recorded activity. The recorded and predicted FRAs for four example units from one animal are shown in the top two rows of Fig. 5a (for results for the second animal, see Supplementary Fig. 5). To visualize the corresponding phenomena at the level of the bottleneck, we applied principal component analysis (PCA) to the bottleneck response *r*^*_b_* and computed the FRAs for the top four components (Fig. 5a, bottom row). While the top component was broadly tuned, the subsequent components appear to provide the building blocks required to create the frequency specificity observed in the output units.

**Fig. 5.**
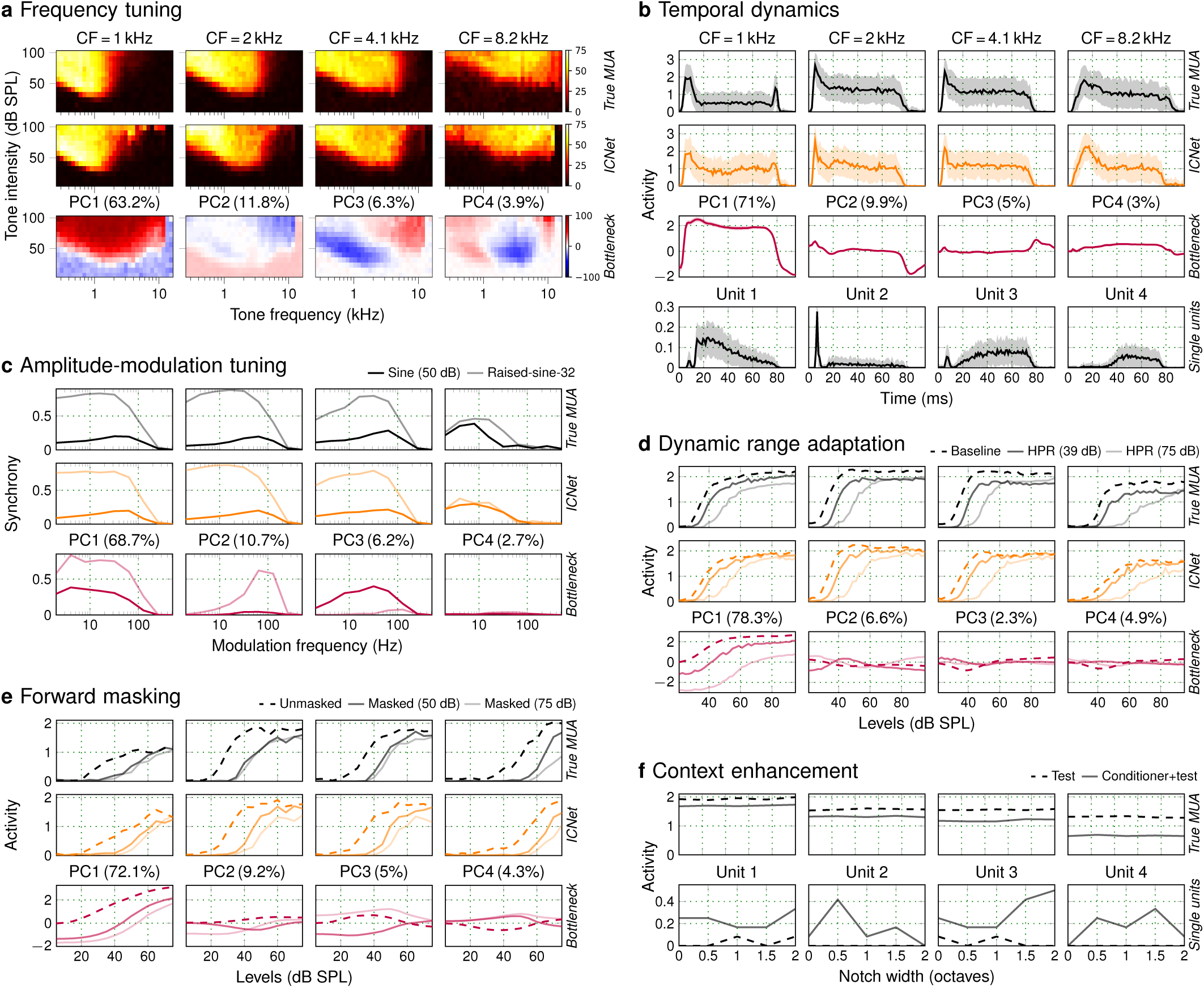
ICNet captures fundamental neurophysiological phenomena. **a.** Recorded (top row) and predicted (middle row) frequency response areas (FRAs) for 4 example neural units with a range of CFs. Each panel shows the overall activity to pure tones of varying intensity and frequency. The FRAs for the top 4 principal components (PCs) of the ICNet bottleneck responses are also shown (bottom row), with the numbers in the parentheses indicating the percentage of the total variance in the bottleneck responses captured by each component. **b.** Temporal dynamics in response to a 2-kHz pure tone stimulus presented at 85 dB SPL. Responses from 4 example single units are also shown (bottom row). **c.** Synchrony to the envelope of an amplitude-modulated narrowband noise (centered at 2 kHz) with two different envelopes. **d.** Rate-intensity functions for broadband noise. The noise intensity was drawn from distributions that were either uniform (baseline) or contained high-probability regions (HPRs) centered at one of two intensities. **e.** Rate-intensity functions for a 2-kHz pure tone that was preceded either by silence (unmasked) or by a masker tone presented at one of two intensities. **f.** Overall activity in response to a multi-tonal complex with a spectral notch centered at 1 kHz. A 1-kHz tone was presented either concurrently with the multi-tonal complex (test) or after some delay (conditioner+test). The overall activity of 4 example single units is also shown (bottom row).

The temporal dynamics of the recorded responses to tones were relatively simple, with a strong onset followed by sustained activity for the duration of the tone. While these dynamics were well captured by ICNet (Fig. 5b, top two rows), it is important to note that the simple dynamics of MUA responses result from the summation of responses across single units that can be more complex, containing, for example, a pause between the onset and sustained components or a build up without a strong onset (Fig. 5b, bottom row). Because ICNet is trained on MUA, it cannot capture these phenomena that are evident only at the level of single units, which may be a limitation for some neurophysiology applications.

To examine tuning for amplitude modulations, we presented tones with two different envelopes – a sinusoid and a raised sinusoid with steeper slopes [26] – and computed the synchrony between the envelope and the neural responses as a function of modulation frequency. ICNet captured the differences in modulation frequency selectivity across units as well as the general increase in synchrony for the raised sinusoid (Fig. 5c, top two rows). At the level of the bottleneck, the top component exhibited low-pass modulation tuning with high synchrony for both envelope types, while the subsequent components were more sharply tuned and selective for a particular envelope type (Fig. 5c, bottom row).

We next examined more complex response properties that are known to be strongly present in the IC. We started with dynamic range adaptation [27], a phenomenon that captures the ability of auditory neurons to adapt their responses to changes in the statistical properties of the acoustic environment. We presented broadband noise with changing intensity that was drawn from distributions that were either uniform or contained high-probability regions (HPRs) centered at particular intensities. We observed clear shifts in dynamic range (illustrated through rate-intensity functions) based on the HPR of the intensity distribution, which were well captured by ICNet and clearly evident in the top component of the bottleneck (Fig. 5d).

We also assessed forward masking [28], a phenomenon in which the neural response to one sound is decreased by the presence of another sound that preceded it. We presented probe tones that were preceded either by silence or by a masker tone presented at different intensities. We observed clear shifts in the rate-intensity functions for the probe tone as a result of the masker, with stronger shifts for the higher-intensity masker. These shifts were again well captured by ICNet and evident in the top component of the bottleneck (Fig. 5e).

Finally, we investigated context enhancement [29], a phenomenon similar to forward masking but with the opposite effect, i.e., an increase in the response to one sound because of the presence of another sound that preceded it. We presented a multi-tonal complex with a spectral notch as context, and then added a tone at the frequency in the center of the notch either concurrently with the multi-tonal complex or after some delay. We found little evidence of context enhancement in the recorded MUA, i.e., responses were decreased rather than increased when the onset of the tone was delayed relative to the context (Fig. 5f, top row). We did, however, observe context enhancement in the responses of some single units (Fig. 5f, bottom row). (Note that this phenomenon is closely related to the temporal dynamics shown in Fig. 5b; units with build-up dynamics are expressing a form of context enhancement.) Overall, ICNet appears to capture many key response properties of IC neurons (excluding those that are evident only at the level of single units), with clear manifestation of the underlying phenomena in its latent dynamics.

## Discussion

In this study, we combined large-scale neural recordings and deep learning to develop ICNet, a comprehensive baseline model of neural coding in the early central auditory pathway. We presented a novel multi-branch encoder-decoder framework to describe the common features of sound encoding across a set of normal-hearing animals, while also accounting for the idiosyncrasy and non-stationarity of individual recordings. ICNet captures the statistical properties of neural activity with high resolution, simulating response patterns across thousands of units with millisecond precision. The simulated responses capture most of the explainable variance in our neural recordings across a wide range of sounds. And because ICNet is based on convolutional operations, it is computationally fast: neural activity can be simulated more than 10x faster than real time, even on a CPU (for a 420 ms sound input, simulation of 512 units takes 49 ms on an Intel Xeon w7-2495X CPU and 39 ms on an Nvidia RTX 4090 GPU).

In order to produce a model that would generalize well, we assembled a broad training dataset that contained representative samples of many complex sounds. We demonstrated that ICNet produced near-perfect simulations of the neural responses to a range of natural and artificial evaluation sounds that were not included in the training set, capturing more than 90% of the explainable variance and correlation in recorded activity for speech in quiet (across all units and animals) and at least 86% of the explainable variance and correlation for speech in noise, music, and broadband moving ripples. ICNet also exhibited key neurophysiological phenomena, such as amplitude modulation tuning and dynamic range adaptation, in both its output units and its latent dynamics. The sounds used to elicit these phenomena (pure tones and noise bursts) are qualitatively different from the complex sounds on which ICNet was trained, and its success in simulating the processing of these sounds is strong evidence of its generality. The primary limitations of ICNet are a reflection of the dataset on which it was trained – MUA from the anesthetized IC – which excludes features of neural responses that are evident only during wakefulness or at the level of single units. Within these constraints, ICNet is highly accurate, particularly for the units with the most reliable responses, and makes few systematic errors.

### ICNet as a baseline model

We designed ICNet with a shared encoder to capture the common features of sound encoding across the individual animals on which it was trained, as most applications of auditory models require a generic simulation of the fundamental features of auditory processing that are common across all normal-hearing individuals. Phenomenological or mechanistic models that are designed by hand will naturally have this property, but models that are learned directly from experimental data will inevitably reflect the idiosyncrasies of the individuals from which the data were collected. The results of our cross-validation experiments (Fig. 3c, Fig. 5 and Supplementary Figs. 4 and 5) suggest that ICNet is generic, i.e., that the inclusion of additional animals during training would not improve its generality.

Because our goal was to develop a baseline model of auditory processing up to the level of the IC, we also designed ICNet so that the effects of non-stationarity (primarily due to accumulated effects of anesthesia over the long duration of the recordings) could be separated from the shared sound encoding and individualized for each animal. By providing ICNet with a second input during training, which reflected the time at which each response was recorded, ICNet was able to capture the non-stationarity with a fixed encoder whose output was modulated as needed at the level of the decoder. After training, ICNet can provide a stationary simulation of neural activity under optimal recording conditions by setting the time input to a fixed value (e.g., 1 hour, corresponding to the beginning of recording). While the non-stationarity in our datasets was a nuisance that we sought to decouple from the shared sound encoding and remove, a similar modeling framework should be capable of capturing non-stationarity that is of interest (e.g., behavioral modulation) given an appropriate dataset for training.

Our use of anesthesia enabled us to collect the large datasets required to train ICNet’s high-capacity encoder. But it is important to note that anesthesia has effects on auditory processing that go beyond the absence of behavioral modulation. The differences between anesthesia and passive wakefulness may be much less pronounced in the IC than in later stages of the auditory pathway, but they are not negligible. These differences are, however, largely quantitative rather than qualitative, i.e., they are restricted to coarse features of neural activity such as overall spike rate or adaptation timescales rather than fine details such as the timing of response events [30]. Thus, it is likely that ICNet’s encoder captures most, if not all, of the features required to simulate activity during passive wakefulness. If simulation of responses during passive wakefulness is needed, this should be achievable with a small dataset that is used to train a relatively simple decoder to map the ICNet bottleneck to the output units.

### ICNet as a foundation core

We demonstrated that ICNet provides a baseline model of early central auditory processing that captures the key features of the mapping from sound to neural activity in the IC. ICNet generalized well across a number of sound classes and could accurately simulate the responses of new animals with just 3 minutes of neural data used to update the decoder weights. There are, of course, many features of auditory processing that ICNet does not capture, such as behavioral modulation or higher-level computations. In this sense, ICNet is analogous to the “core” module in the foundation model of visual processing that was developed by Wang et al. [31]. ICNet could serve a similar role in models of auditory processing, providing a baseline representation that can be modulated based on behavioral factors or further processed to reflect cortical computations. The use of ICNet as a foundation core should significantly reduce the amount of data required to develop the modulatory mappings or updated decoders for advanced models.

Another critical area for further development is binaural processing. We designed ICNet to capture the mapping between a single sensory input (a diotic stimulus in which the same sound was presented in both ears) and the neural activity recorded from both hemispheres of the IC. But IC activity is strongly modulated by binaural disparity, and these modulations encode information that is critical for spatial hearing. Fortunately, the features of neural activity that encode binaural disparity are largely separable from those that encode other acoustic information [32], so it is likely that spatial hearing can also be captured using a module that modulates the baseline representation provided by ICNet.

### Potential applications of ICNet

For hearing research, ICNet opens new possibilities with potential benefits that are both scientific (studies are no longer data limited) and ethical (animal experiments can be limited to confirmatory studies). Much of hearing research is focused on understanding the cortical mechanisms that allow for stream segregation and recognition of objects in complex auditory scenes. Understanding these mechanisms is impossible without first specifying precisely the representation of the auditory scene that is made available to the cortex for further analysis. ICNet has the potential to be a valuable tool in this regard, allowing downstream computations to be referenced directly to early neural representations. Cochlear models are already employed widely and successfully in this way [33, 34]; using ICNet as a second reference point midway along the central auditory pathway would enable early and late central processing to be decoupled.

ICNet also provides a comprehensive baseline normal-hearing reference for technology development. It can be used as a front-end in the design of systems that achieve natural (rather than objectively optimal) performance, as its outputs reflect the physiological constraints that are faced by the brain. We demonstrated that ICNet’s bottleneck response provides a compact representation of sound encoding that can be effective as a front-end for speech recognition and can be used on its own for applications that require a general auditory model, rather than explicit neural simulation. The use of ICNet for optimization of signal processing (e.g., noise suppression) could unlock additional perceptual benefit beyond that achieved through optimization based directly on acoustics. The same may also be true for the development of assistive listening technologies, with the output of ICNet serving as a target reference against which the efficacy of different hearing restoration strategies can be compared.

## Methods

### Experimental protocol

Experiments were performed on 14 young-adult gerbils of both sexes that were born and raised in standard laboratory conditions. IC recordings were made at an age of 20-28 weeks. All experimental protocols were approved by the UK Home Office (PPL P56840C21).

### Preparation for IC recordings

Recordings were made using the same procedures as in previous studies [21, 22]. Animals were placed in a sound-attenuated chamber and anesthetized for surgery with an initial injection of 1 ml per 100 g body weight of ketamine (100 mg per ml), xylazine (20 mg per ml), and saline in a ratio of 5:1:19. The same solution was infused continuously during recording at a rate of approximately 2.2 µl per min. Internal temperature was monitored and maintained at 38.7^◦^ C. A small metal rod was mounted on the skull and used to secure the head of the animal in a stereotaxic device. The pinnae were removed and speakers (Etymotic ER-10X) coupled to tubes were inserted into both ear canals. Sounds were low-pass filtered at 12 kHz (except for tones) and presented at 44.1 kHz without any filtering to compensate for speaker properties or ear canal acoustics. Two craniotomies were made along with incisions in the dura mater, and a 256-channel multi-electrode array was inserted into the central nucleus of the IC in each hemisphere.

### Multi-unit activity

Neural activity was recorded at 20 kHz. MUA was measured from recordings on each channel of the electrode array as follows: (1) a bandpass filter was applied with cutoff frequencies of 700 and 5000 Hz; (2) the standard deviation of the background noise in the bandpass-filtered signal was estimated as the median absolute deviation / 0.6745 (this estimate is more robust to outlier values, e.g., neural spikes, than direct calculation); (3) times at which the bandpass filtered signal made a positive crossing of a threshold of 3.5 standard deviations were identified and grouped into 1.3 ms bins.

### Characteristic frequency analysis

To determine the preferred frequency of the recorded units, 50 ms tones were presented with frequencies ranging from 294 Hz to 16384 Hz in 0.2 octave steps (without any high-pass filtering) and intensities ranging from 4 dB SPL to 85 dB SPL in 9 dB steps. Tones were presented 8 times each in random order with 10 ms cosine on and off ramps and 75 ms pause between tones. The characteristic frequency (CF) of each unit was defined as the frequency that elicited a significant response at the lowest intensity. Significance was defined as *p <* 0.0001 for the total MUA count elicited by a tone given an assumed Poisson distribution with rate equal to the higher of either the mean count observed in time windows of the same duration in the absence of sound or 1/8.

### Single-unit activity

Single-unit spikes were isolated using Kilosort as described in [22]. Recordings were separated into overlapping 1-hour segments with a new segment starting every 15 minutes. Kilosort 1 [35] was run separately on each segment and clusters from separate segments were chained together. Clusters were retained for analysis only if they were present for at least 2.5 hours of continuous recording.

### Deep neural network models

#### Time-invariant single-branch architecture

The time-invariant single-branch DNN that was used to predict the neural activity of individual recordings is shown in Fig. 1. The model was trained to map a sound waveform *s* to neural activity *R*^^^ via an inferred conditional distribution of MUA counts *p*(*R^*|*s*) using the following architecture: (1) a SincNet layer [36] with 48 bandpass filters of size 64 and stride of 1, followed by a symmetric logarithmic activation *y* = sgn(*x*) ∗ log(|*x*| + 1); (2) a stack of five 1-D convolutional layers with 128 filters of size 64 and stride of 2, each followed by a PReLU activation; (3) a 1-D bottleneck convolutional layer with 64 filters of size 64 and a stride of 1, followed by a PReLU activation; and (4) a decoder convolutional layer without bias and 512 × *N_c_* filters of size 1, where *N_c_* is the number of parameters in the count distribution *p*(*R^*|*s*) as described below. All convolutional layers in the encoder included a bias term and used a causal kernel. A cropping layer was added after the bottleneck to remove the left context from the output, which eliminated any convolutional edge effects.

For the Poisson model, *N_c_* = 1 and the decoder output was followed by a softplus activation to produce the rate function *λ* that defined the conditional count distribution for each unit in each time bin. (Exponential activation was also tested, but softplus yielded better performance.) The loss function was 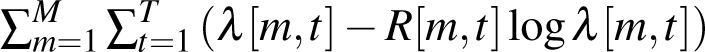, where *M* and *T* are the number of units and time bins in the response, respectively. For the crossentropy model, *N_c_*= 5 and the decoder output was followed by a softmax activation to produce the probability of each possible count *n* ∈ {0*,…, N_c_* − 1} for each unit in each time bin. The loss function was 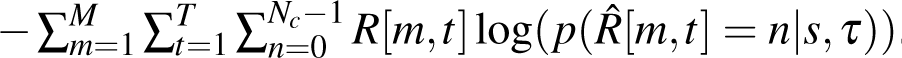. The choice of 4 as the maximum possible count was made based on the MUA count distributions across all datasets (the percentage of counts above 4 was less than 0.02%). All neural activity was clipped during training and inference at a maximum value of 4.

#### Time-variant single-branch architecture

The time-variant single-branch architecture (Figs. 2-3) included an additional time input *τ*. Each sample of the time input defined the relative time at which the corresponding sound sample was presented during a given recording (range of approximately 0-12 hr). To match the dimensionality and dynamic range of the sound input after the SincNet layer, the time input was given to a convolutional layer with 48 filters of size 1 (without bias), followed by a symmetric logarithmic activation. The resulting output was then concatenated to the output of the SincNet layer to form the input of the first convolutional layer of the encoder. The loss function was 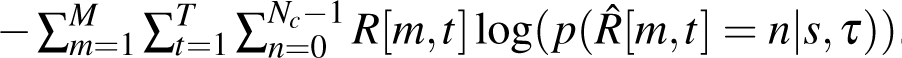.

#### Time-variant multi-branch architecture

The final ICNet model (Figs. 3-4) comprises a time-variant DNN with a shared encoder and separate decoder branches for each of the individual animals. The architecture of the encoder and decoders were identical to those in the single-branch models, except for the additional time input. Instead of a single time input to the encoder, a separate time input was used for each animal with the same dimensionality and sampling rate as the bottleneck output. Each sample of the time input thus defined the relative time at which a corresponding MUA sample was recorded, and was given to a convolutional layer with 64 filters of size 1 (with bias), followed by a PReLU activation. The output of each of these layers was then multiplied (elementwise) by the bottleneck output and was given to the corresponding decoder.

For the additional analysis in Supplementary Fig. 3, a time-variant multi-branch architecture with a more complex time module was trained. Each time input was given to a cascade of 4 convolutional layers with 128 filters of size 1 (and bias), followed by PReLU activations. The final output of each time module was multiplied by the bottleneck output and given to the corresponding decoder as in the final ICNet.

#### Linear-nonlinear models

For the comparison of ICNet with linear-nonlinear models in Fig. 3d, single-layer models for seven animals were trained through Poisson regression. The models comprised a convolutional layer with 512 filters (one for each output unit) of size 64 samples and stride 1 (without bias), followed by an exponential activation. Mel spectrograms were used as inputs and were computed as follows: (1) the short-time Fourier transform (STFT) of the audio inputs was computed using a window of 512 samples and a step size of 32 samples, resulting in spectrograms with time resolution of 762.9395 Hz (to match the 1.3ms time bins of the neural activity); (2) the power spectrograms (squared magnitude of the STFT outputs) were mapped to 96 Mel bands with frequencies logarithmically spaced between 50 Hz to 12 kHz; (3) the logarithm of the Mel spectrograms was computed and was offset by 7 dB so that the spectrograms comprised only positive values; (4) a sigmoid activation function was applied to normalize the range of the spectrograms. An L2 regularization penalty was additionally applied to the convolutional kernel during training using a factor of 10^-5^. A number of parameters for the spectrogram inputs (STFT window size, linear or Mel scale, number of frequency bands, activation functions) were tested and the values that produced the best performance were used.

### Training

Models were trained to transform 24414.0625 Hz sound input frames of 8192 samples into 762.9395 Hz neural activity frames of 256 samples (Fig. 1b). Context of 2048 samples was added on the left side of the sound input frame (total of 10240 samples) and was cropped after the bottleneck layer (2048 divided by a decimation factor of 2^5^ resulted in 64 cropped samples at the level of the bottleneck). Sound inputs were scaled such that an RMS of 0.04 corresponded to a level of 94 dB SPL. For the time-variant models, time inputs were expressed in seconds and were scaled by 1/36000 to match the dynamic range to that of the sound inputs. These scalings were necessary to enforce training with sufficiently high decimal resolution, while maximally retaining the datasets’ statistical mean close to 0 and standard deviation close to 1 to accelerate training.

The DNN models were trained on NVidia RTX 4090 GPUs using Python and Tensorflow [37]. A batch size of 50 was used with the Adam optimizer and a starting learning rate of 0.0004. All trainings were performed so that the learning rate was halved if the loss in the validation set did not decrease for 2 consecutive epochs. Early stopping was used and determined the total number of training epochs if the validation loss did not decrease for 4 consecutive epochs. This resulted in an average of 38.1 ± 3.4 epochs for the nine single-branch models, and a total of 42 epochs for the nine-branch ICNet model. To speed up training, the weights from one of the single-branch models were used to initialize the encoder of the multi-branch models.

### Training dataset

All sounds that were used for training the DNN models totaled to 7.83 hr and are described below. 10% of the sounds were randomly chosen to form the validation set during training. The order of the sound presentation was randomized for each animal during recording. Multi-branch models were trained using a dataset of 7.05 hr, as some of the music sounds were not recorded across all nine animals.

#### Speech

Sentences were taken from the TIMIT corpus [38] that contains speech read by a wide range of American English talkers. The entire corpus excluding “SA” sentences was used and was presented either in quiet or with background noise. The intensity for each sentence was chosen at random from 45, 55, 65, 75, 85, or 95 dB SPL. The speech-to-noise ratio (SNR) was chosen at random from either 0 or 10 when the speech intensity was 55 or 65 dB SPL (as is typical of a quiet setting such as a home or an office) or –10 or 0 when the speech intensity was 75 or 85 dB SPL (as is typical of a noisy setting such as a pub). The total duration of speech in quiet was 1.25 hr and the total duration of speech in noise was 1.58 hr.

#### Noise

Background noise sounds were taken from the Microsoft Scalable Noisy Speech Dataset [39], which includes recordings of environmental sounds from a large number of different settings (e.g., café, office, roadside) and specific noises (e.g., washer-dryer, copy machine, public address announcements). The intensity of the noise presented with each sentence was determined by the intensity of the speech and the SNR as described above.

#### Processed speech

Speech from the TIMIT corpus was also processed in several ways: (1) the speed was increased by a factor of 2 (via simple resampling without pitch correction of any other additional processing); (2) linear multi-channel amplification was applied, with channels centered at 0.5, 1, 2, 4, and 8 kHz and gains of 3, 10, 17, 22, and 25 dB SPL, respectively; or (3) the speed was increased and linear amplification was applied. The total duration of processed speech was 2.08 hr.

#### Music

Pop music was taken from the musdb18 dataset [40], which contains music in full mixed form as well as in stem form with isolated tracks for drums, bass, vocals and other (e.g., guitar, keyboard). The total duration of the music presented from this dataset was 1.28 hr. A sub-portion of this music was used when training multi-branch models, which corresponded to 0.67 hr. Classical music was taken from the musopen dataset (https://musopen.org; including piano, violin and orchestral pieces) and was presented either in its original form; after its speed was increased by a factor of 2 or 3; after it was high-pass filtered with a cutoff frequency of 6 kHz; or after its speed was increased and it was high-pass filtered. The total duration of the music presented from this dataset was 0.8 hr.

#### Moving ripples

Dynamic moving ripple sounds were created by modulating a series of sustained sinusoids to achieve a desired distribution of instantaneous amplitude and frequency modulations. The lowest frequency sinusoid was either 300 Hz, 4.7 kHz, or 6.2 kHz. The highest frequency sinusoid was always 10.8 kHz. The series contained sinusoids at frequencies between the lowest and the highest in steps of 0.02 octaves, with the phase of each sinusoid chosen randomly from between 0 and 2*π*. The modulation envelope was designed so that the instantaneous frequency modulations ranged from 0 to 4 cycles/octave, the instantaneous amplitude modulations ranged from 0 to 10 Hz, and the modulation depth was 50 dB. The total duration of the ripples was 0.67 hr.

### Abridged training dataset

For the analysis in Fig. 5 (and Supplementary Figs. 5 and 4), a sub-portion of all sounds that were described above were used. The abridged training dataset was generated by keeping only the first 180 s of each speech segment (60% of total duration), while preserving the total duration of the remaining sounds. The resulting dataset totaled to 5.78 hr (347 minutes) and included 2.95 hr of (processed and unprocessed) speech and 2.83 hr of music and ripples. As before, 10% of the sounds were randomly chosen to form the validation set during training.

### Evaluation

To predict neural activity using the trained DNN models, the decoder parameters were used to define either a Poisson distribution or a categorical probability distribution with *N_c_*classes using the Tensorflow Probability toolbox [41]. The distribution was then sampled from to yield simulated neural activity across time bins and units, as shown in Fig. 1b.

#### Evaluation dataset

All model evaluations used only sounds that were not part of the training dataset. Each sound segment was 30 s in duration. The timing of the presentation of the evaluation sounds varied across animals. For two animals, each sound was presented only twice and the presentation times were random. For three animals, each sound was presented four times, twice in successive trials during the first half of the recording and twice in successive trials during the second half of the recording. For the other four animals, each sound was presented ten times, twice in successive trials at times that were approximately 20, 35, 50, 65, and 80% of the total recording time, respectively.

#### Speech in quiet

For all animals, a speech segment from the UCL SCRIBE dataset (http://www.phon.ucl.ac.uk/resource/scribe) consisting of sentences spoken by a male talker was presented at 60 dB SPL. For four of the animals (Fig. 4b), another speech segment from the same dataset was chosen and was presented at 85 dB SPL, consisting of sentences spoken by a female talker.

#### Speech in noise

For all animals, a speech segment from the UCL SCRIBE dataset consisting of sentences spoken by a female talker was presented at 85 dB SPL in hallway noise from the Microsoft Scalable Noisy Speech dataset at 0 dB SNR. For four of the animals (Fig. 4b), another speech segment from the same dataset was chosen and was presented at 75 dB SPL, consisting of sentences spoken by a male talker mixed with café noise from the Microsoft Scalable Noisy Speech dataset at 4 dB SNR.

#### Moving ripples

For all animals, dynamic moving ripples with the lowest frequency sinusoid of 4.7 kHz were presented at 85 dB SPL. For four of the animals (Fig. 4b), dynamic moving ripples with the lowest frequency sinusoid of 300 Hz were also used and were presented at 60 dB SPL.

#### Music

For all animals, three seconds from each of 10 mixed pop songs from the musdb18 dataset were presented at 75 dB SPL. For four of the animals (Fig. 4b), a solo violin recording from the musopen dataset was presented at 85 dB SPL.

#### Pure tones

For measuring Fano factors (Fig. 1) and assessing systematic errors (Supplementary Fig. 2b), 75 ms tones at 4 frequencies (891.44, 2048, 3565.78 and 8192 Hz) were presented at either 59 or 85 dB SPL with 10 ms cosine on and off ramps. Each tone was repeated 128 times, and the resulting responses were used to compute the average, the variance and the full distribution of MUA counts (using time bins from 2.5 ms to 80 ms).

### Neurophysiological evaluation dataset

For the analysis in Fig. 5 (and Supplementary Figs. 4 and 5), a dataset was recorded that included sounds commonly used in experimental studies to assess the neurophysiological properties of IC neurons (frequency tuning, temporal dynamics, rate-intensity functions, dynamic range adaptation, forward masking and context enhancement). This evaluation dataset was presented to two animals that were not included in ICNet training, interspersed with sounds from the abridged training dataset described above.

Principal component analysis (PCA) was used to visualize the manifestation of the relevant phenomena in the latent representation of the ICNet bottleneck. PCA was performed separately for each sound, after subtracting the average activity of each bottleneck channel to silence and dividing the resulting responses by -500 (to match the scale and sign of the MUA for plotting).

#### Frequency tuning

For the results shown in Fig. 5a, Supplementary Fig. 5a and Supplementary Fig. 4e, tones were presented at frequencies from 256 to 16384 Hz (1/5 octave spacing) and intensities from 4 to 103 dB SPL (9 dB spacing) with 50 ms duration and 10 ms cosine on and off ramps. Each tone was presented 8 times in random order with 75 ms of silence between presentations. The responses were used to compute frequency response area (FRA) heatmaps by averaging activity across time bins from 7.9 ms to 50 ms after tone onset.

#### Temporal dynamics

For the results shown in Fig. 5b and Supplementary Fig. 5b, tones were presented at frequencies from 294.1 to 14263.1 Hz (1/5 octave spacing) and intensities of 59 and 85 dB SPL with 75 ms duration and 10 ms cosine on and off ramps. Each tone was presented 128 times in random order with 75 ms of silence between presentations. The responses were used to compute the average and standard deviation of MUA counts in time bins from 0 to 82.9 ms after tone onset.

#### Dynamic range adaptation

For the results shown in Fig. 5d and Supplementary Fig. 5d, the sounds defined in [27] were used. For the baseline rate-intensity functions, noise bursts were presented at intensities from 21 to 96 dB SPL (2 dB spacing) with 50 ms duration. Each intensity was presented 32 times in random order with 300 ms of silence between presentations. For the rate-intensity functions of high-probability region (HPR) sounds, noise bursts were presented at intensities from 21 to 96 dB SPL (1 dB spacing) with 50 ms duration. The intensities were randomly drawn from a distribution with an HPR of 12 dB range centered at either 39 or 75 dB SPL to create sequences of 16.15 s (no silence was added between noise bursts). Each value in the HPR was drawn 20 times and all other values drawn once. Sixteen different random sequences were presented. The responses were used to compute the rate-intensity functions by averaging activity across time bins from 7.9 ms to 50 ms after noise onset.

#### Amplitude-modulation tuning

For the results shown in Fig. 5c, Supplementary Fig. 5c, and Supplementary Fig. 4g-h, the sounds defined in [26] were used. Noise was generated with a duration of 1 s (with 4 ms cosine on and off ramps) and a bandwidth of one octave, centered at frequencies from 500 to 8000 Hz (1/2 octave spacing). The noise was either unmodulated or modulated with 100% modulation depth at frequencies from 2 to 512 Hz (1 octave spacing). Two different envelope types were used for modulating the noise: a standard sinusoid and a sinusoid raised to the power of 32 (raised-sine-32 envelope in Fig. 1A of [26]). All sounds were presented 6 times in random order with 800 ms of silence between presentations. In an attempt to present the sounds at approximately 30 dB above threshold for most units, each sound was presented at two intensities that varied with center frequency, with higher intensities used for very low and very high frequencies to account for the higher thresholds at these frequencies. The baseline intensities were 50 and 65 dB SPL with the addition of [15 0 0 0 20 20] dB SPL for center frequencies of [500, 1000, 2000, 4000, 8000] Hz. The additional intensities for other center frequencies were determined by interpolating between these values.

To obtain modulation transfer functions, the fast Fourier transform of the responses was computed using time bins from 7.9 ms to 1007.9 ms after sound onset with a matching number of frequency bins (763 bins). Synchrony was defined as the ratio between the modulation frequency component and the DC component (0 Hz) of the magnitude-squared spectrum for all modulation frequencies.

#### Forward masking

For the results shown in Fig. 5e, Supplementary Fig. 5e and Supplementary Fig. 4f, the sounds defined in [4] were used. For the unmasked rate-intensity functions, tones (probes) were presented at frequencies from 500 and 11313.71 Hz (1/2 octave spacing) with 20 ms duration and 10 ms cosine on and off ramps. The baseline intensities of the tones varied from 5 and 75 dB SPL (5 dB spacing) and the intensity was increased for very low and very high frequencies as described above. Each tone was presented 12 times in random order with 480 ms of silence between tones.

For the masked rate-intensity functions, a masking tone at the same frequency as the probe was presented with a duration of 200 ms and 10 ms cosine on and off ramps, with a 10 ms pause added between the offset of the masker and the onset of the probe. The masker was presented at baseline intensities of 50 and 75 dB SPL and the intensity was increased for very low and very high frequencies as for the probe tone, but with the additional intensities scaled by 2/3. The responses were used to compute the rate-intensity functions by averaging actvity across time bins from 7.9 ms to 20 ms after probe onset.

#### Context enhancement

For the results shown in Fig. 5g and Supplementary Fig. 5g, the sounds defined in [29] were used. The conditioner and test sounds were generated by adding together equal-amplitude tonal components with random phases and frequencies from 200 Hz to 16 kHz (1/10 octave spacing). A notch was carved out of the spectrum with center frequency from 500 to 8000 Hz (1/2 octave spacing) and width from 0 to 2 octaves (0.5 octave spacing). For the test sound, an additional tone was added at the center frequency of the notch. The conditioner and test sounds had durations of 500 ms and 100 ms, respectively, with 10 ms cosine on and off ramps. The sounds were presented at baseline intensities of 40 and 60 dB SPL and the intensity was increased for very low and very high frequencies as described above. Each test sound was presented either alone or preceded by the conditioner 12 times in random order with 1.2 s of silence between presentations. The responses were used to compute the average activity across time bins from 7.9 ms to 100 ms after test sound onset.

##### Non-stationarity

For Fig. 2g, we quantified the non-stationarity in each recording as the absolute difference of the covariance of neural activity elicited by the same sound on two successive trials and on two trials recorded several hours apart. The covariance was computed after flattening (collapsing neural responses across time and units into one dimension). To obtain a single value for the covariance on successive trials, the values from the two pairs of successive trials (one pair from early in the recording and one pair from late in the recording) were averaged. To obtain a single value for the covariance on separated trials, the values from the two pairs of separated trials were averaged.

##### Overall performance metrics

RMSE and log-likelihood were used as overall performance metrics (Figs. 1-3). The reported results for both metrics were the averages computed over two trials of each sound within each individual recording. One trial was chosen from the first half of the recording and the other from the second, resulting in times of 3.51 ± 1.83 hr and 8.92 ± 1.16 hr across all nine animals. For each trial, the RMSE and log-likelihood were computed as follows:

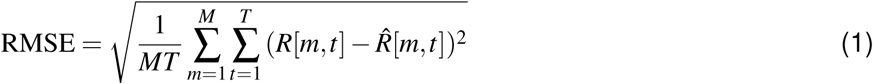

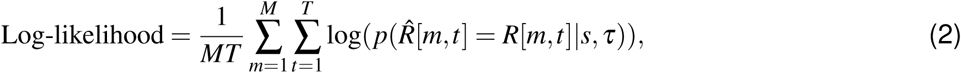

where *M* is the number of units, *T* is the number of time bins, *s* is the sound stimulus, *τ* is the time input, *R* is the recorded neural activity, and *R^* is the predicted neural activity obtained by sampling from the inferred distribution *p*(*R^*|*s, τ*) (or *p*(*R^*|*s*) for a time-invariant model).

##### Predictive power metrics

To assess the predictive power of ICNet (Figs. 2 and 4 and Supplementary Figs. 3 and 4), we selected two metrics that were computed using successive trials for the seven animals for which this was possible (chosen from the beginning of the recording, resulting in times of 1.94 ± 0.46 hr across animals). The two metrics are formulated as the fraction of explainable variance or correlation that the model explained across all units. The fraction of explainable correlation explained was computed as follows:

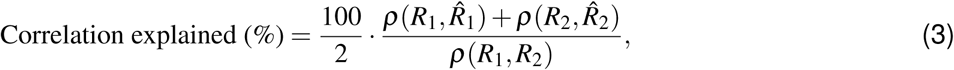

where *R*_1_*, R*_2_ ∈ N*^M^*^×*T*^ are the recorded neural activity for each trial, *R^*_1_*, R*^^^_2_ ∈ N*^M^*^×*T*^ are the predicted neural activity, and *ρ* is the correlation coefficient. The responses were flattened (collapsed across time and units into one dimension) before the correlation was computed.

The fraction of explainable variance explained was computed as follows [14]:

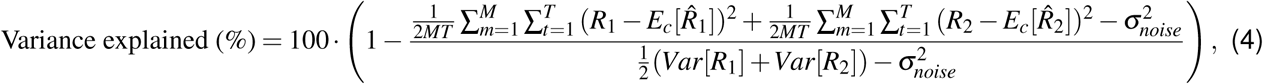

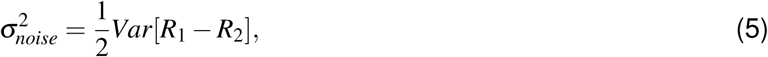

where *E_c_* denotes expectation over counts of *p*(*R^*|*s, τ*) (or *p*(*R^*|*s*) for a time-invariant model). The responses were flattened (collapsed across time and units into one dimension) before the variances in the denominator of Eq *4* and in Eq *5* were computed, while flattening was achieved in the numerator of Eq *4* through the averaging across units and time. To compute the predictive power of ICNet for each unit (Fig. 4d), we computed the formulas of Eqs *3*-*5* across time only (without collapsing across the unit dimension or averaging across units). Values above 100% for individual units indicate that the ICNet responses were more similar to the recorded responses than the recorded responses were to each other across successive trials, which is possible in recordings with high non-stationarity.

To visualize coherence in Supplementary Fig. 2a, we computed the magnitude-squared coherence for recorded and predicted responses to the four primary evaluation sounds. The coherence of the recorded responses across trials was computed as follows:

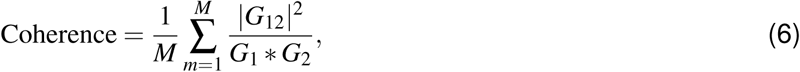

where *M* is the number of units, *G*_12_ is the cross-spectral density between the responses *R*_1_*, R*_2_ for the two successive trials and *G*_1_*, G*_2_ the power spectral density for each of the responses, respectively. The spectral densities were computed across the first dimension of the neural responses (time) using 763 frequency bins and a Hanning window. The coherence of the predicted and recorded responses was obtained by computing the coherence between the predicted and recorded response on each trial using the same formula and taking the average of the two values.

#### Phoneme recognition

A DNN architecture based on Conv-TasNet [42] was used to train an automatic-speech-recognition (ASR) back-end that predicted phonemes from the ICNet bottleneck response. The DNN architecture comprised: (1) a convolutional layer with 64 filters of size 3 and no bias, followed by a PReLU activation, (2) a normalization layer, followed by a convolutional layer with 128 filters of size 1, (3) a block of 8 dilated convolutional layers (dilation from 1 to 2^7^) with 128 filters of size 3, including PReLU activations and residual skip connections in between, (4) a convolutional layer with 256 filters of size 1, followed by a sigmoid activation, (5) an output convolutional layer with 40 filters of size 3, followed by a softmax activation. All convolutional layers were 1-D and used a causal kernel with a stride of 1.

The ASR model was trained using the train subset of the TIMIT corpus [38]. The TIMIT sentences were resampled to 24414.0625 Hz and were segmented into sound input frames of 81920 samples (windows of 65536 samples with 50% overlap and left context of 16384 samples). In each training step, the sound input frames were given to the frozen ICNet model to generate bottleneck responses at 762.9395 Hz (output frames of 2048 samples with left context of 256 samples). The bottleneck response was then given to the ASR back-end to predict the probabilities of the 40 phoneme classes across time (output frames of 2048 samples at 762.9395 Hz after cropping the context).

The TIMIT sentences were calibrated to randomly selected levels between 40 and 90 dB SPL in steps of 5 dB, and were randomly mixed with the 18 noise types from the DEMAND dataset [43] at SNRs of -30, -20, -10, 0, 10, 20, 30, and 100 dB. The phonetic transcriptions of the TIMIT dataset were downsampled to the sampling frequency of the neural activity (762.9395 Hz) and were grouped into 40 classes (15 vowels and 24 consonants plus the glottal stop). To account for the phonetic class imbalance (high prevalence of silence in the dataset), a focal crossentropy loss function was used for training with the focusing parameter *γ* set to 5. ASR models were trained to predict speech recognition from the ICNet bottleneck response, as well as from the bottleneck responses of the nine single-branch time-variant models. The trained ASR models were evaluated using 6 random sentences of the test subset of the TIMIT corpus. The sentences were calibrated at 65 dB SPL, and speech-shaped noise was generated to match the long-term average spectrum of the sentences. A baseline system was also trained with the same DNN architecture, but using a standard Mel spectrogram as the input. The Mel spectrogram was computed from the sound input (24414.0625 Hz) using frames of 512 samples, a hop size of 32 samples and 64 Mel channels with frequencies logarithmically spaced from 50 to 12000 Hz. These parameters were chosen to maximize performance while matching the resolution of the ICNet bottleneck response (762.9395 Hz output frames with 64 channels).

### Statistics

Confidence intervals for all reported values were computed using 1000 bootstrap samples. For the metrics that we used to assess model performance (RMSE, log-likelihood, variance and correlation explained), bootstrapping was performed across units for each animal. For the ASR results (Fig. 3f), bootstrapping was performed across animals.

## Data availability

All of the sounds and IC recordings that were used to evaluate ICNet in Fig. 4b and d are available via https://doi.org/10.5281/zenodo.12636772. The four sounds can be used as inputs to ICNet to simulate responses for comparison with the IC recordings provided for seven animals. Researchers seeking access to the full set of neural recordings for research purposes should contact the corresponding author via e-mail to set up a material transfer agreement.

## Code availability

The code of the ICNet model is available via https://doi.org/10.5281/zenodo.11943090 or https://github.com/fotisdr/ICNet. A Jupyter notebook is included with a simple usage example for ICNet. A non-commercial, academic UCL license applies.

## Acknowledgments

The authors thank Jennifer Bizley for her advice.

## Author contributions

**F.D.**: Conceptualization, Data Curation, Formal analysis, Investigation, Methodology, Software, Validation, Visualization, Writing: Original Draft Preparation; **L.P.**: Conceptualization, Methodology, Software; **S.S.**: Investigation; **Y.X.**: Investigation; **A.F.**: Conceptualization, Project administration, Software, Supervision; **N.A.L.**: Conceptualization, Data Curation, Funding acquisition, Investigation, Methodology, Project Administration, Resources, Supervision, Writing - Review & Editing

## Supplementary information

**Supplementary Fig. 1.**
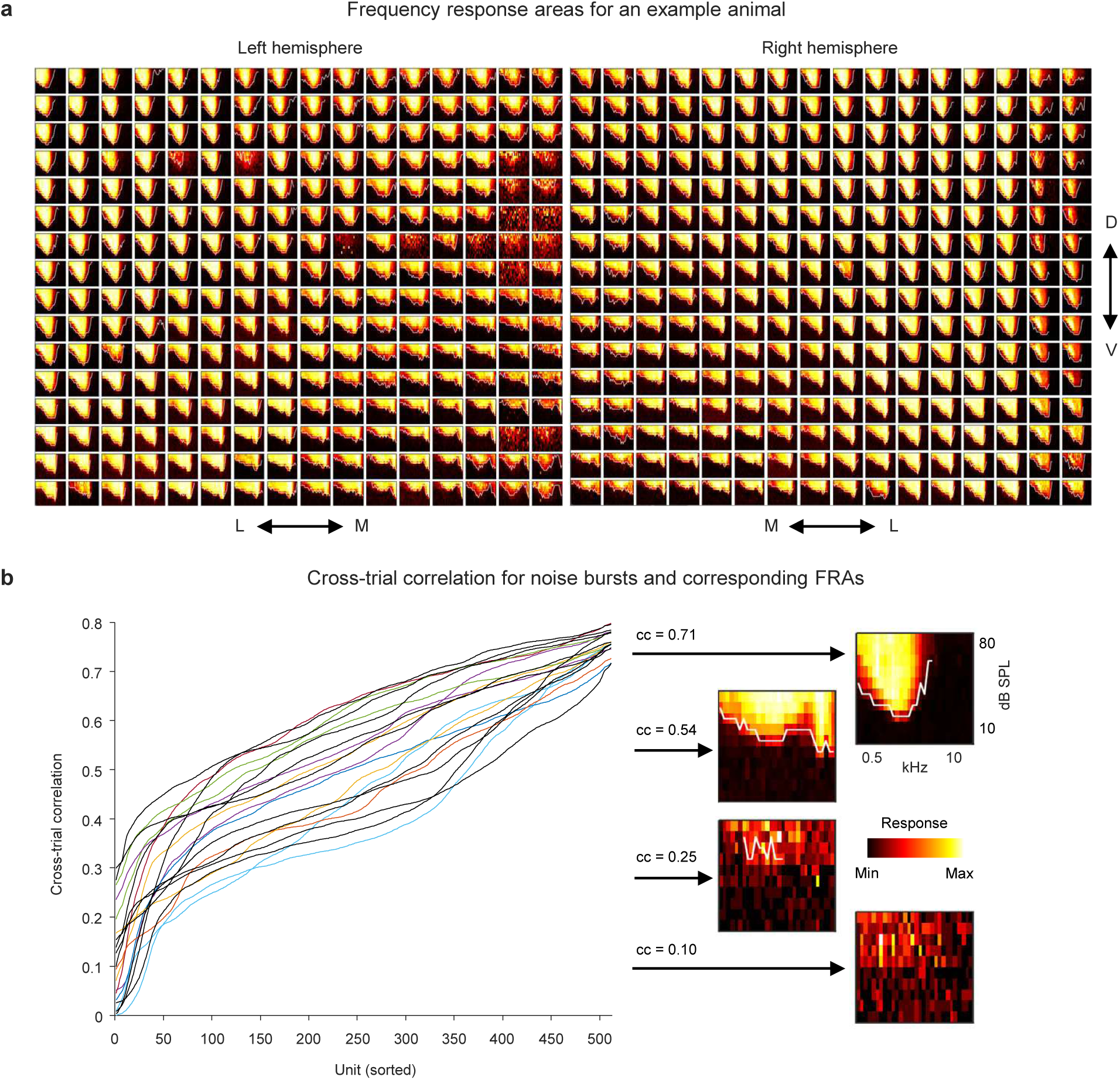
Identification of “good” recording sites. We have described our approach to targeting the central nucleus of the inferior colliculus in a previous paper (see Fig. S1 in [22]). We used the same approach for the recordings that we used in the development of ICNet. Our electrodes are designed to span the mediolateral extent of the central nucleus. Panel **a** shows frequency response areas (FRAs) estimated from MUA recorded on 512 recording sites for one example animal (each group of two columns represents 1 of 8 shanks on each electrode; the arrows indicate medial, lateral, dorsal, and ventral). Aside from a few scattered damaged sites, most of the sites exhibit the clear “V-shaped” FRAs associated with the central nucleus. Only some of the most medial sites on the left electrode and the most lateral sites on the right electrode exhibit FRAs that are not clear and V-shaped. As an objective quantitative criterion to identify (undamaged) sites that are likely to lie within the central nucleus, we calculated the cross-trial correlation of responses to broadband noise bursts and then sorted the sites by this cross-trial correlation value in increasing order. The results after sorting are shown in panel **b** for 20 normal hearing animals, with the 9 animals used to train ICNet shown in black and other animals shown in color. Sites with high cross-trial correlation values also have clear V-shaped FRAs, whereas sites with low cross-trial correlation values do not. We included all sites in model training but excluded sites with cross-trial correlation *<* 0.2 for evaluation.

**Supplementary Fig. 2.**
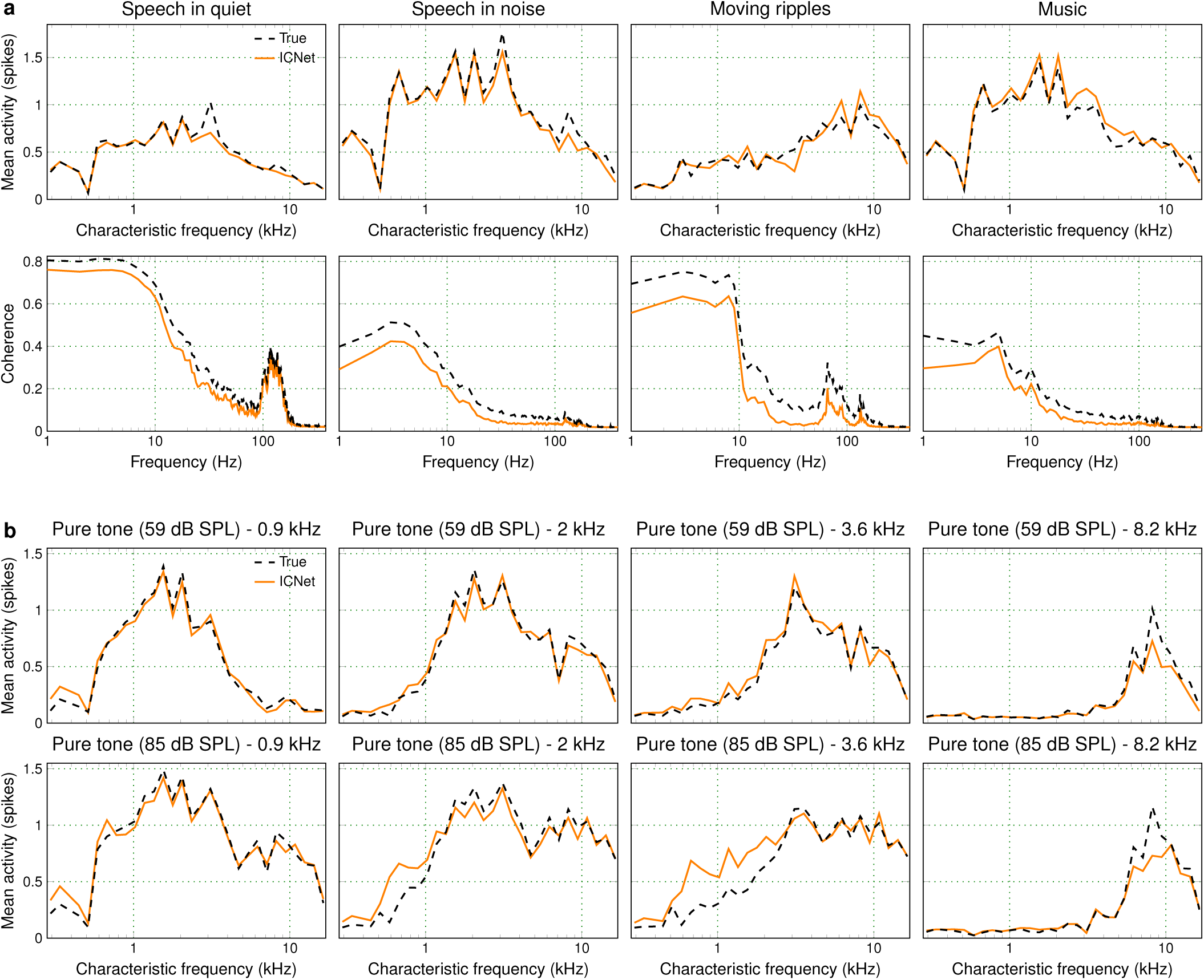
Assessment of systematic errors. The top row of panel **a** shows the overall activity (mean of counts across time bins) of 3476 units from 7 animals along with that predicted by ICNet in response to our 4 primary evaluation sounds. The results are grouped and sorted based on the CF of each unit. The bottom row of panel **a** shows the average coherence spectrum (across units) between either the recorded responses across trials or the recorded and predicted responses for the same 4 sounds. Overall, we found little evidence of systematic errors in ICNet’s predictions of responses to complex sounds. Panel **b** shows the overall activity of 4446 units from 9 animals along with that predicted by ICNet in response to pure tones at 4 frequencies and 2 intensities. When presented with pure tones of 2 and 3.6 kHz at 85 dB SPL, ICNet overpredicted the overall activity for units with CFs below the tone frequency (frequencies between 0.5 and 2 kHz). This was not evident for the tones presented at 59 dB SPL.

**Supplementary Fig. 3.**
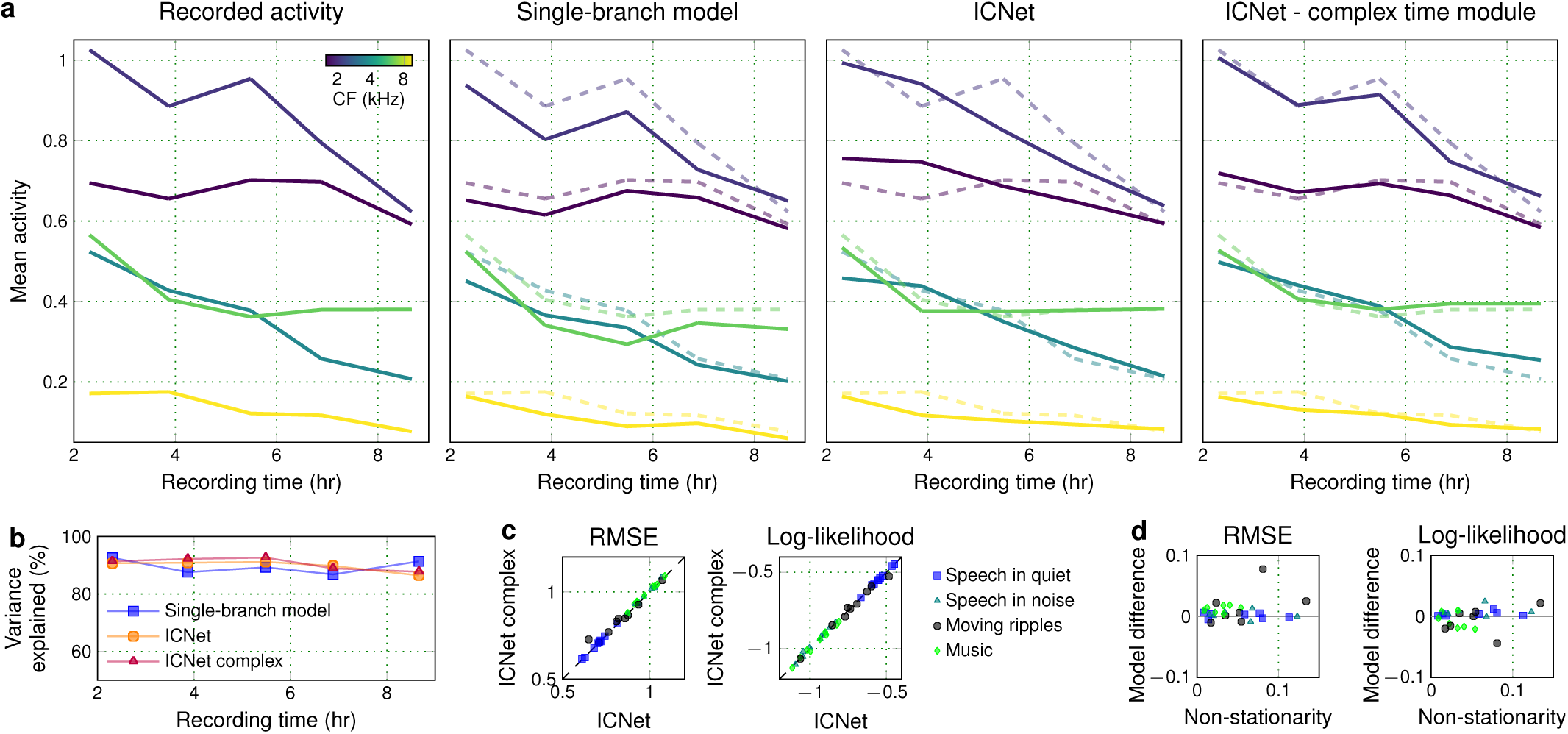
Accounting for non-stationarity in neural recordings. The long duration of our recordings had varying effects across units, with non-uniform and, in some cases, non-monotonic changes in overall activity for non-stationary recordings. We tested time modules with varying capacity in the ICNet architecture, but found that the additional capacity required to capture the full complexity of non-stationarity did not significantly improve overall performance. The first plot in panel **a** shows the mean activity of 5 example units (from the non-stationary recording of Fig. 2b) in response to 5 presentations of the same speech sound at different times, with the units chosen to reflect the range of non-uniform and non-monotonic effects of non-stationarity in our recordings. The second plot in panel **a** compares the recorded activity to the predictions of the single-branch time-variant architecture (Fig. 2), which was able to capture the full effects of the non-stationarity. The third plot in panel **a** shows the predictions of the final ICNet, which uses a time module with limited capacity and is only able to capture monotonic trends. The fourth plot in panel **a** shows the predictions of ICNet with a more complex time module (see Methods), which was again able to capture the full effects of the non-stationarity. We compared the performance of the 3 model architectures (single-branch, final ICNet, and ICNet with complex time module) with respect to their overall predictive power as in Fig. 2. Panel **b** shows the median predictive power of the 3 model architectures across all units at 5 different times during the non-stationary recording of Fig. 2b. Panel **c** shows a performance comparison across 9 animals and 4 sounds between the two variations of ICNet. Panel **d** shows the performance difference between the two varations of ICNet (ICNet with complex time module − final ICNet) as a function of recording non-stationarity across 7 animals and 4 sounds. Because there was little evidence of performance benefit for the more complex time module, we opted for the simpler time module for the final ICNet.

**Supplementary Fig. 4.**
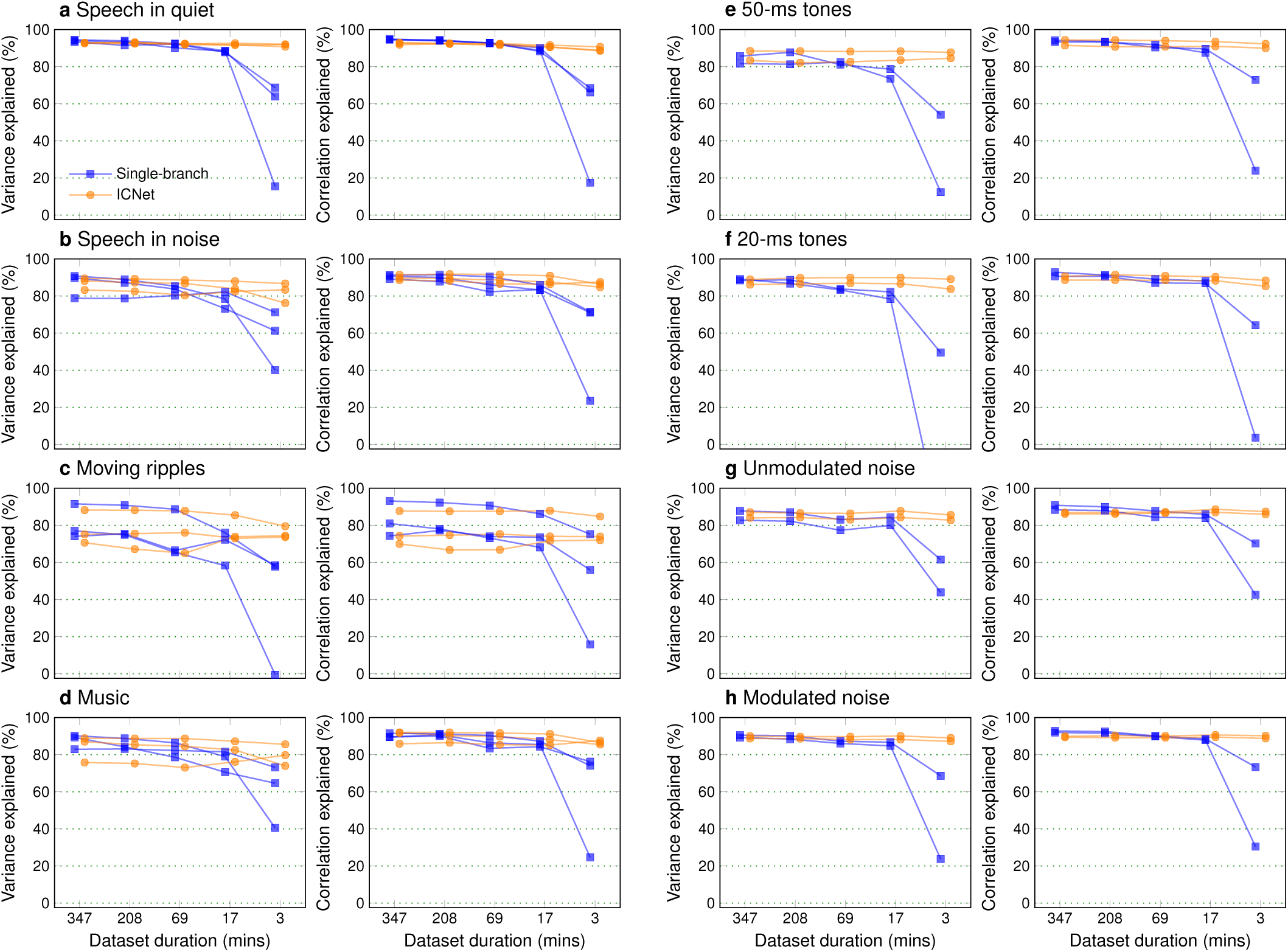
Generality of ICNet across new sounds and animals. We used neural recordings from new animals (unseen during ICNet training) to train single-branch and ICNet-based models. We trained the single-branch models from scratch (learning all parameters), and trained the ICNet-based models by freezing the ICNet encoder and learning only a new decoder. We trained both model variants using the abridged training dataset (347 minutes; see Methods) and sub-portions of it which were generated by keeping 60%, 20%, 5% and 1% of the dataset, resulting in training datasets with durations of 208, 69, 17 and 3 minutes, respectively. We then assessed the performance of all trained models across different sounds (unseen during training) as a function of the amount of data used for training. Panels **a**-**d** show model performance across 3 new animals on the 4 sounds that were used for our primary ICNet evaluation (Fig. 4). Panels **e**-**h** show model performance across 2 new animals on the sounds that were used to assess frequency tuning, forward masking and amplitude-modulation tuning (Fig. 5; sounds from the neurophysiological evaluation dataset for which we recorded responses to repeated trials). The results show that (1) ICNet performed as well as fully-trained single-branch models and (2) that this level of performance could be achieved with as little as 3 minutes of data to retrain the decoder.

**Supplementary Fig. 5.**
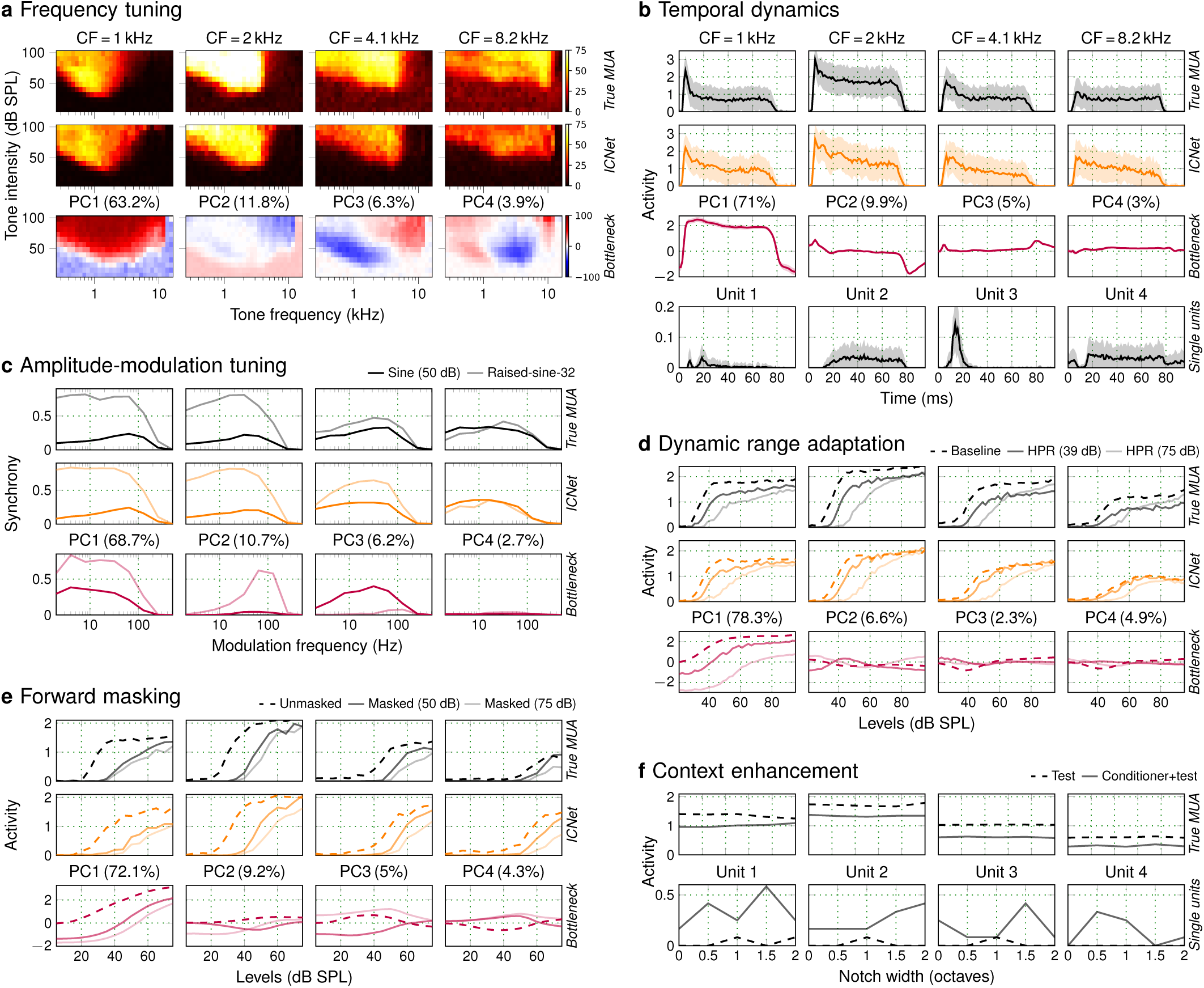
ICNet captures fundamental neurophysiological phenomena. The results of Fig. 5, repeated for another animal.

## References

1. Osses Vecchi A, Varnet L, Carney LH, Dau T, Bruce IC, Verhulst S, et al. A comparative study of eight human auditory models of monaural processing. Acta Acustica. 2022;6:17.

2. Meyer AF, Williamson RS, Linden JF, Sahani M. Models of neuronal stimulus-response functions: elaboration, estimation, and evaluation. Frontiers in Systems Neuroscience. 2017;10:109.

3. Rahman M, Willmore BD, King AJ, Harper NS. Simple transformations capture auditory input to cortex. Proceedings of the National Academy of Sciences. 2020;117(45):28442–28451.

4. Nelson PC, Carney LH. A phenomenological model of peripheral and central neural responses to amplitude-modulated tones. The Journal of the Acoustical Society of America. 2004;116(4):2173– 2186.

5. Chi T, Ru P, Shamma SA. Multiresolution spectrotemporal analysis of complex sounds. The Journal of the Acoustical Society of America. 2005;118(2):887–906.

6. Carney LH, Li T, McDonough JM. Speech coding in the brain: representation of vowel formants by midbrain neurons tuned to sound fluctuations. eNeuro. 2015;2(4).

7. Farhadi A, Jennings SG, Strickland EA, Carney LH. Subcortical auditory model including efferent dynamic gain control with inputs from cochlear nucleus and inferior colliculus. The Journal of the Acoustical Society of America. 2023;154(6):3644–3659.

8. Baby D, Van Den Broucke A, Verhulst S. A convolutional neural-network model of human cochlear mechanics and filter tuning for real-time applications. Nature Machine Intelligence. 2021;3(2):134–143.

9. Drakopoulos F, Baby D, Verhulst S. A convolutional neural-network framework for modelling auditory sensory cells and synapses. Communications Biology. 2021;4(1):827.

10. Nagathil A, Bruce IC. WaveNet-based approximation of a cochlear filtering and hair cell transduction model. The Journal of the Acoustical Society of America. 2023;154(1):191–202.

11. Van Den Broucke A, Drakopoulos F, Baby D, Verhulst S. Otoacoustic emissions in a deep-neural-network model of cochlear mechanics. In: The 14th International Mechanics of Hearing Workshop (MoH 2022). vol. 3062; 2024.

12. McIntosh L, Maheswaranathan N, Nayebi A, Ganguli S, Baccus S. Deep learning models of the retinal response to natural scenes. Advances in Neural Information Processing Systems. 2016;29.

13. Walker EY, Sinz FH, Cobos E, Muhammad T, Froudarakis E, Fahey PG, et al. Inception loops discover what excites neurons most using deep predictive models. Nature Neuroscience. 2019;22(12):2060– 2065.

14. Cadena SA, Denfield GH, Walker EY, Gatys LA, Tolias AS, Bethge M, et al. Deep convolutional models improve predictions of macaque V1 responses to natural images. PLoS Computational Biology. 2019;15(4):e1006897.

15. Lurz KK, Bashiri M, Willeke K, Jagadish AK, Wang E, Walker EY, et al. Generalization in data-driven models of primary visual cortex. bioRxiv. 2020; p. 2020.10.05.326256.

16. Bashivan P, Kar K, DiCarlo JJ. Neural population control via deep image synthesis. Science. 2019;364(6439):eaav9436.

17. Kell AJ, Yamins DL, Shook EN, Norman-Haignere SV, McDermott JH. A task-optimized neural network replicates human auditory behavior, predicts brain responses, and reveals a cortical processing hierarchy. Neuron. 2018;98(3):630–644.

18. Keshishian M, Akbari H, Khalighinejad B, Herrero JL, Mehta AD, Mesgarani N. Estimating and interpreting nonlinear receptive field of sensory neural responses with deep neural network models. eLife. 2020;9:e53445.

19. Pennington JR, David SV. A convolutional neural network provides a generalizable model of natural sound coding by neural populations in auditory cortex. PLoS Computational Biology. 2023;19(5):e1011110.

20. Garcia-Lazaro JA, Belliveau LA, Lesica NA. Independent population coding of speech with submillisecond precision. Journal of Neuroscience. 2013;33(49):19362–19372.

21. Sabesan S, Fragner A, Bench C, Drakopoulos F, Lesica NA. Large-scale electrophysiology and deep learning reveal distorted neural signal dynamics after hearing loss. eLife. 2023;12:e85108.

22. Armstrong AG, Lam CC, Sabesan S, Lesica NA. Compression and amplification algorithms in hearing aids impair the selectivity of neural responses to speech. Nature Biomedical Engineering. 2022;6(6):717–730.

23. Lyamzin DR, Garcia-Lazaro JA, Lesica NA. Analysis and modelling of variability and covariability of population spike trains across multiple time scales. Network: Computation in Neural Systems. 2012;23(1-2):76–103.

24. Pecka M, Siveke I, Grothe B, Lesica NA. Enhancement of ITD coding within the initial stages of the auditory pathway. Journal of Neurophysiology. 2010;103(1):38–46.

25. Williamson RS, Sahani M, Pillow JW. The equivalence of information-theoretic and likelihood-based methods for neural dimensionality reduction. PLoS Computational Biology. 2015;11(4):e1004141.

26. Kim DO, Carney L, Kuwada S. Amplitude modulation transfer functions reveal opposing populations within both the inferior colliculus and medial geniculate body. Journal of Neurophysiology. 2020;124(4):1198–1215.

27. Dean I, Harper NS, McAlpine D. Neural population coding of sound level adapts to stimulus statistics. Nature Neuroscience. 2005;8(12):1684–1689.

28. Nelson PC, Smith ZM, Young ED. Wide-dynamic-range forward suppression in marmoset inferior colliculus neurons is generated centrally and accounts for perceptual masking. Journal of Neuroscience. 2009;29(8):2553–2562.

29. Nelson PC, Young ED. Neural correlates of context-dependent perceptual enhancement in the inferior colliculus. Journal of Neuroscience. 2010;30(19):6577–6587.

30. Ter-Mikaelian M, Sanes DH, Semple MN. Transformation of temporal properties between auditory midbrain and cortex in the awake Mongolian gerbil. Journal of Neuroscience. 2007;27(23):6091–6102.

31. Wang EY, Fahey PG, Ding Z, Papadopoulos S, Ponder K, Weis MA, et al. Foundation model of neural activity predicts response to new stimulus types. Nature. 2025;640(8058):470–477.

32. Belliveau LA, Lyamzin DR, Lesica NA. The neural representation of interaural time differences in gerbils is transformed from midbrain to cortex. Journal of Neuroscience. 2014;34(50):16796–16808.

33. Shan T, Cappelloni MS, Maddox RK. Subcortical responses to music and speech are alike while cortical responses diverge. Scientific Reports. 2024;14(1):789.

34. Saddler MR, McDermott JH. Models optimized for real-world tasks reveal the necessity of precise temporal coding in hearing. bioRxiv. 2024; p. 2024.04.21.590435.

35. Pachitariu M, Steinmetz N, Kadir S, Carandini M, Kenneth D H. Kilosort: realtime spike-sorting for extracellular electrophysiology with hundreds of channels. BioRxiv. 2016; p. 061481.

36. Ravanelli M, Bengio Y. Speaker recognition from raw waveform with SincNet. In: 2018 IEEE Spoken Language Technology Workshop (SLT); 2018. p. 1021–1028.

37. Abadi M, Barham P, Chen J, Chen Z, Davis A, Dean J, et al. Tensorflow: A system for large-scale machine learning. In: 12th USENIX Symposium on Operating Systems Design and Implementation (OSDI 16); 2016. p. 265–283.

38. Garofolo JS, Lamel LF, Fisher WM, Fiscus JG, Pallett DS. DARPA TIMIT acoustic-phonetic continous speech corpus CD-ROM. NIST speech disc 1-1.1. NASA STI/Recon technical report n. 1993;93:27403.

39. Reddy CK, Beyrami E, Pool J, Cutler R, Srinivasan S, Gehrke J. A scalable noisy speech dataset and online subjective test framework. arXiv. 2019; p. 1909.08050.

40. Rafii Z, Liutkus A, Stöter FR, Mimilakis SI, Bittner R. The MUSDB18 corpus for music separation; 2017.

41. Dillon JV, Langmore I, Tran D, Brevdo E, Vasudevan S, Moore D, et al. Tensorflow distributions. arXiv. 2017; p. 1711.10604.

42. Luo Y, Mesgarani N. Conv-TasNet: Surpassing Ideal Time–Frequency Magnitude Masking for Speech Separation. IEEE/ACM Transactions on Audio, Speech, and Language Processing. 2019;27(8):1256–1266.

43. Thiemann J, Ito N, Vincent E. The diverse environments multi-channel acoustic noise database (DEMAND): A database of multichannel environmental noise recordings. In: Proceedings of Meetings on Acoustics. vol. 19; 2013.

